# Mammalian SWI/SNF complex activity regulates POU2F3 and constitutes a targetable dependency in small cell lung cancer

**DOI:** 10.1101/2024.01.21.576304

**Authors:** Leslie Duplaquet, Kevin So, Alexander W. Ying, Xinyue Li, Yixiang Li, Xintao Qiu, Rong Li, Shilpa Singh, Xiaoli S. Wu, Qi Liu, Jun Qi, Tim D.D. Somerville, Hillary Heiling, Emanuele Mazzola, Yenarae Lee, Thomas Zoller, Christopher R. Vakoc, John G. Doench, William C. Forrester, Tinya Abrams, Henry W. Long, Matthew J. Niederst, Cigall Kadoch, Matthew G. Oser

## Abstract

Small cell lung cancers (SCLC) are comprised of heterogeneous subtypes marked by lineage-specific transcription factors, including ASCL1, NEUROD1, and POU2F3. POU2F3-positive SCLC, ∼12% of all cases, are uniquely dependent on POU2F3 itself; as such, approaches to attenuate POU2F3 expression may represent new therapeutic opportunities. Here using genome-scale screens for regulators of POU2F3 expression and SCLC proliferation, we define mSWI/SNF complexes, including non-canonical BAF (ncBAF) complexes, as top dependencies specific to POU2F3-positive SCLC. Notably, clinical-grade pharmacologic mSWI/SNF inhibition attenuates proliferation of all POU2F3-positive SCLCs, while disruption of ncBAF via BRD9 degradation is uniquely effective in pure non-neuroendocrine POU2F3-SCLCs. mSWI/SNF maintains accessibility over gene loci central to POU2F3-mediated gene regulatory networks. Finally, chemical targeting of SMARCA4/2 mSWI/SNF ATPases and BRD9 decrease POU2F3-SCLC tumor growth and increase survival *in vivo*. Taken together, these results characterize mSWI/SNF-mediated global governance of the POU2F3 oncogenic program and suggest mSWI/SNF inhibition as a therapeutic strategy for SCLC.

## Introduction

Small cell lung cancer (SCLC) is a high-grade neuroendocrine cancer that accounts for ∼15% of lung cancers^1–3^. Nearly all SCLCs are genetically driven by loss-of-function (LOF) mutations in tumor suppressor genes *RB1* and *TP53*^4–6^ without recurrent mutations in druggable oncogenic drivers. SCLC are heterogeneous and broadly consist of four molecular subtypes, including the neuroendocrine subtypes marked by ASCL1 (∼60%) and NEUROD1 (∼20%), and the non-neuroendocrine subtypes marked by the POU-family transcription factor POU2F3 (∼10-12%) or the inflammatory subtype (∼10%)^3,7–10^. Notably, the molecular subtypes marked by high expression of lineage-specific transcription factors (TFs) ASCL1, NEUROD1, and POU2F3 offer potential therapeutic opportunities as these TFs represent selective dependencies in the SCLCs expressing them^11–16^. In particular, the POU2F3 SCLC subtype is highly dependent on expression of POU2F3 itself or the recently discovered POU2F3 co-activators OCA-T1 (gene names *c11ORF53*/*POU2AF2*) or OCA-T2 (gene names *COLCA2*/*POU2AF3*)^13–16^.

The POU2F3 (or OCT-11) TF assembles as an octamer around the ATGCAAAT DNA motif^17^. SCLCs that express POU2F3 are associated with low expression of neuroendocrine markers including INSM1, synaptophysin, and chromogranin A^7^. Apart from SCLC and neuroendocrine prostate cancer^18^, expression of POU2F3 is restricted to and essential for the development of tuft cells, mucosal epithelial cells found in various epithelial tissues such as the airway, nasopharyngeal, and the intestine^19^. Indeed, POU2F3-knockout mice are completely viable and only exhibit loss of tuft cells^20–23^. Despite this exquisite cell specificity and likely limited resulting toxicity if targeted, chemical disruption of POU2F3 itself and its transcriptional co-activators OCA-T1 and OCA-T2 remain an unmet challenge. With this, a major goal in the field has been to define druggable regulators of POU2F3 expression or gene regulatory function as an opportunity for new therapeutic strategies in POU2F3-positive SCLC.

Here we use a positive selection CRISPR/Cas9-based screening strategy to identify genes that when inactivated decrease POU2F3 expression in SCLC cellular models. We uncover selective dependencies on the mammalian SWI/SNF (mSWI/SNF or BAF) family of ATP-dependent chromatin remodeling complexes, large multi-subunit assemblies that govern genomic accessibility and gene expression^24^ and which are frequently mutated in human cancers^25^. An emerging body of evidence implicates mSWI/SNF complexes in the maintenance and oncogenicity of a range of tumor types^26–33^ which has prompted the development of small molecule inhibitors and degraders that are currently being evaluated in clinical trials^34–36^(<u>NCT04879017</u>, <u>NCT04891757</u>). We define the mechanisms by which mSWI/SNF remodelers regulate POU2F3 activity at the chromatin and gene regulatory levels and demonstrate that pharmacologic disruption of mSWI/SNF ATPase paralog subunits, SMARCA4 and SMARCA2, as well as the non-canonical BAF (ncBAF) subunit, BRD9, using clinical-grade inhibitors FHD-286 and FHD-609, respectively, dramatically attenuate SCLC growth in culture and in vivo. These data expand the growing repertoire of cancers in which mSWI/SNF complexes are leveraged to orchestrate oncogenic gene expression and suggest new therapeutic opportunities for aggressive POU2F3-positive SCLC.

## Results

### Genome-scale positive selection screen identifies key POU2F3 regulators

We recently developed a positive selection-based screening approach to identify regulators of an oncogene of interest by fusing the oncogene to a modified version of deoxycytidine kinase (oncogene-DCK*)^37^. In this strategy, BVdU selectively kills cells that express oncogene-DCK* while cells that downregulate oncogene-DCK* are BVdU-resistant and hence survive BVdU treatment (**Figure 1A**). This assay was previously optimized for small molecule and CRISPR/Cas9 screens with initial iterations of the assay involving lentiviral overexpression of an oncogene-DCK* under the control of a ubiquitous promoter (e.g. CMV) in either HEK-293T or Jurkat cells^37^, and hence did not report on endogenous gene regulation. Here we modified the DCK*/BVdU CRISPR/Cas9 screening approach to identify regulators of endogenous expression of *POU2F3*, a lineage-specific TF oncogene demarcating 10-15% of SCLCs^3,7,14^.To do this, we used CRISPR/Cas9 to knock-in DCK*, a self-cleaving peptide (P2A), and GFP to create an POU2F3-DCK*-P2A-GFP fusion in the NCI-H1048 SCLC cell line that expresses and is dependent on POU2F3 (**Figure 1B**)^13,14^. Two rounds of FACS sorting for GFP-positive cells were performed to obtain a pure population of NCI-H1048 POU2F3-DCK*-P2A-GFP cells (**Figure 1C, S1A**). As expected, NCI-H1048 POU2F3-DCK*-P2A-GFP cells were highly sensitive to BVdU relative to NCI-H1048 parental cells with a large therapeutic window at a BVdU concentration of 10 μM, indicative of screening suitability (**Figure 1D**). Further, CRISPR-mediated inactivation of DCK reduced BVdU sensitivity, demonstrating BVdU anti-proliferative impact was due to DCK* expression (**Figure S1B,C**).

**Fig. 1.**
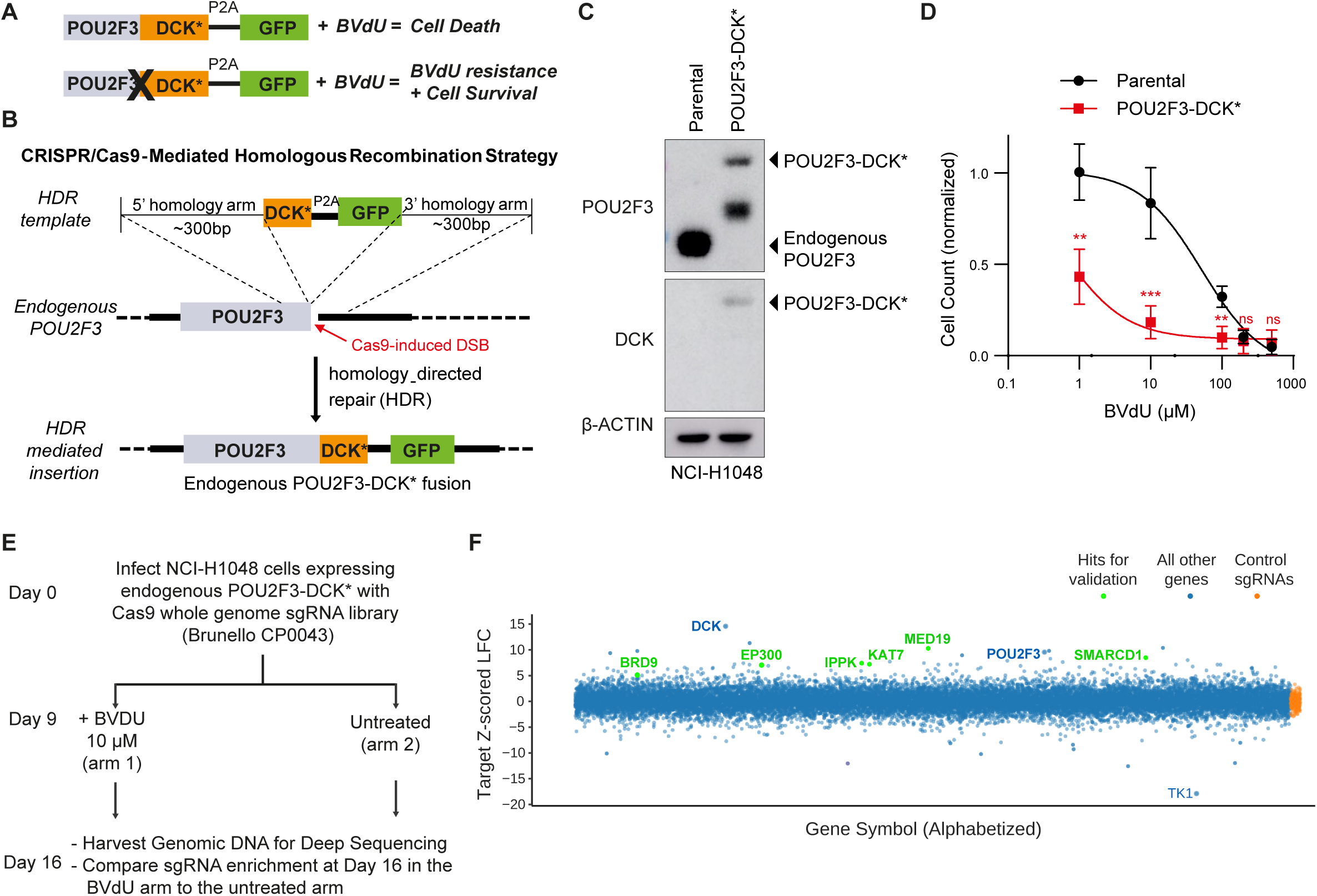
Genome-scale positive selection screen identifies key POU2F3 regulators. (**A**) Schema for positive selection assay to unbiasedly identify regulators of POU2F3 where DCK* variant allows the incorporation of the cytotoxic BVdU metabolites into newly synthesized DNA which are otherwise not toxic to wild-type cells. DCK*=variant of the deoxycytidine kinase. BVdU=BromoVinyl deoxyUridine. (**B**) Schema of the CRIS-PR/Cas9-mediated homologous recombination strategy used to knock-in the suicide gene DCK* and GFP (linked by P2A) into the endogenous POU2F3 locus of the human NCI-H1048 SCLC cell line. These “knock-in” (KI) cells endogenously express POU2F3-DCK*-P2A-GFP chimeric fusion protein. (**C**) Immunoblot analysis of NCI-H1048 engineered to endogenously express POU2F3-DCK* using the strategy in B compared to NCI-H1048 parental cells. (**D**) Dose response assays with BVdU of the indicated cell lines from C treated with BVdU for 7 days showing viable cell counts relative to the DMSO control. n = 4 biological replicates. **=p<0.01, ***-p<0.001 using a 2-tailed unpaired t-test. error bars represent mean +/− SEM. (**E**) Schema of the CRISPR/Cas9 positive selection screen performed in NCI-H1048 POU2F3-DCK*-P2A-GFP cells infected with the whole genome Brunello sgRNA library. Following selection, cells were treated with BVdU (10µM) or untreated for 7 days and then harvested to extract genomic DNA for deep sequencing. (**F**) Apron analysis from the positive-selection CRISPR/Cas9 screen in E comparing BVdU treated vs. untreated NCI-H1048 POU2F3-DCK*-P2A-GFP cells. n=2 biological replicates. Enriched hits for validation are marked in green.

To identify endogenous regulators of POU2F3, we next subjected NCI-H1048 POU2F3-DCK*-P2A-GFP cells to CRISPR-Cas9 resistance screens using the genome-wide Brunello sgRNA library containing 77,741 targeting sgRNAs and 1000 control sgRNAs at an MOI of 0.5^38^ (**Figure 1E**). Following transduction and selection, cells were maintained in culture, and on Day 9, all cells were pooled and an early timepoint was harvested. Remaining cells were split into 2 arms (10 μM BVdU or DMSO), maintaining representation of at least 500 cells/sgRNA in each arm and grown in either BVdU or DMSO for 7 days (Day 16) at which point the cells were harvested, genomic DNA was isolated, and deep sequencing was performed.

Analysis of changes in sgRNA abundance identified several sgRNAs exhibiting robust enrichment in both biological replicates in BVdU-versus untreated samples, demonstrating technical success of the screen (**Figures S1D-F, Table S1**). Importantly, among the top-scoring guides were those corresponding to *DCK* and *POU2F3* (**Figure S1F**). Notably, gene level analyses comparing BVdU and untreated conditions at the Day 16 time point highlighted several top-scoring sgRNAs with scores at or near those of *DCK* and *POU2F*3, including those targeting chromatin regulatory genes *MED19, EP300,* and *KAT7*, signaling pathway members such as *IPPK*, and several members of the mammalian SWI/SNF (BAF) family of ATP-dependent chromatin remodeling complexes, particularly the non-canonical (ncBAF) complex^39^ including *SMARCD1* and *BRD9* (**Figure 1F, Table S1**).

### Non-canonical BAF (ncBAF) complexes regulate POU2F3 expression and proliferation in POU2F3-positive SCLC

To validate our findings, we first performed CRISPR-based inactivation of several highly significant top-enriched hits. Guides targeting *SMARCD1*, *EP300*, *MED19*, *IPPK*, and *KAT7*, all markedly reduced total POU2F3 protein levels as well as cellular proliferation in the NCI-H1048 SCLC cell line (**Figure 2A, S2A, Table S1**). We then sought to determine which enriched hits are selective dependencies for POU2F3-positive SCLC relative to other SCLC cell lines and all other cancer cell lines. To do this, we curated and analyzed recent DepMap (Project Achilles, Broad Institute) datasets (23Q2 release) reporting on genome-scale CRISPR-Cas9-based gene knockout screening performed across >1000 cancer cell lines to date^40^. 23 cell lines of SCLC origin are included in this dataset, including all four POU2F3-positive cell lines (NCI-H1048, NCI-H211, NCI-H526 and COR-L311). Strikingly we found that POU2F3-positive SCLC lines were significantly dependent on ncBAF components *SMARCD1* and *BRD9* as well as the conserved mSWI/SNF ATPase subunit, SMARCA4, relative to other SCLC cell lines and all other cancer cell lines profiled (**Figure 2B,C, S2B-F**). Further, in contrast to mSWI/SNF components, POU2F3 dependency did not significantly correlate with the dependencies of other hits validated to impact POU2F3 expression, including *EP300*, *MED19*, *IPPK*, and *KAT7* (**Figure S2C-G**).

**Fig. 2.**
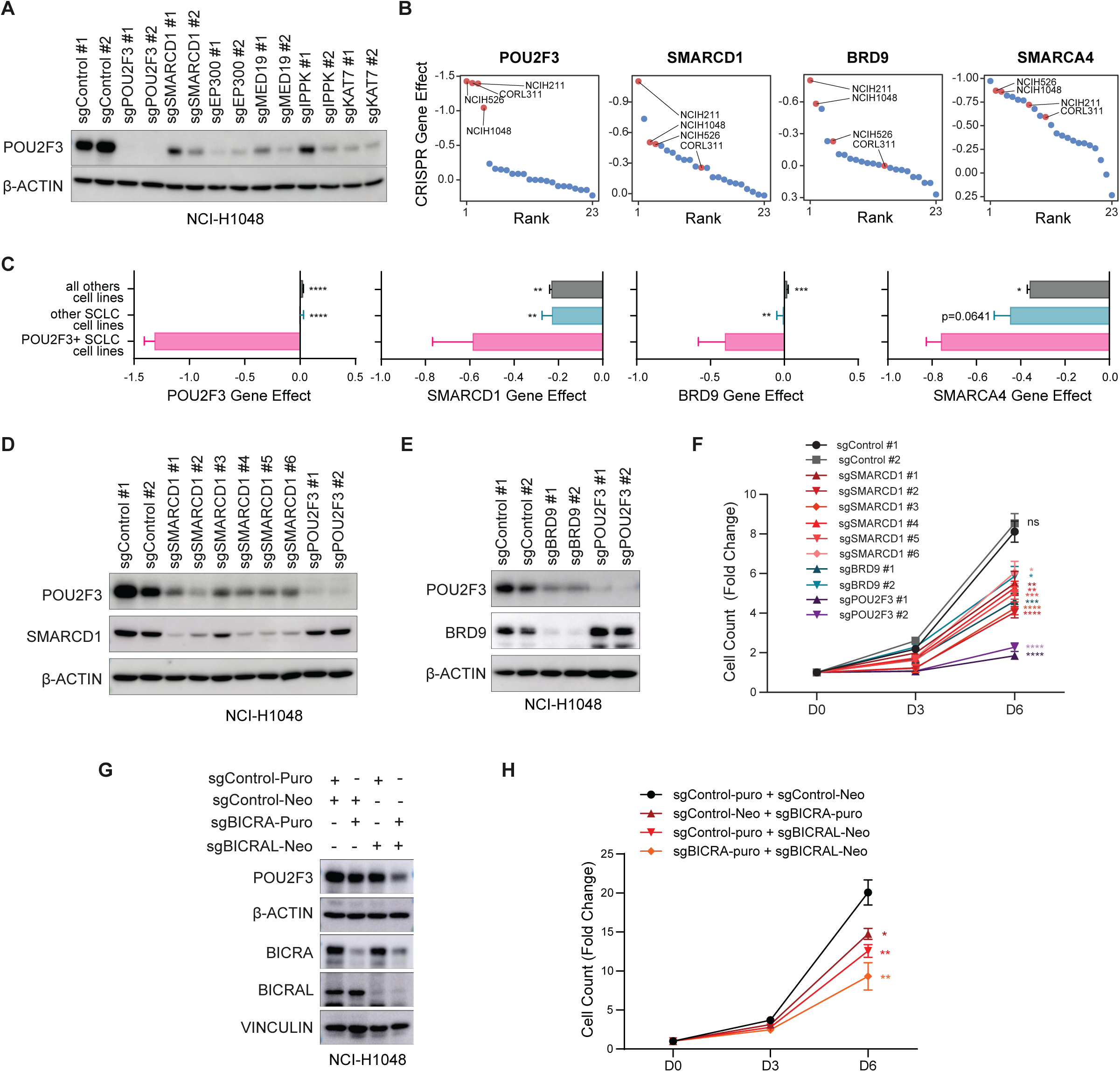
Non-canonical BAF (ncBAF) complexes regulate POU2F3 expression and proliferation in POU2F3-positive SCLC. (**A**) Immunoblot analysis of parental NCI-H1048 cells transduced with sgRNAs targeting genes that scored as candidate POU2F3 regulators or non-targeting controls (sgControl). Immunoblotting was performed 6 days after transduction and is representative of 4 biological replicates. (**B**) Cell line rank vs. CRISPR gene effect for POU2F3, SMARCD1, BRD9, and SMARCA4 of the 23 SCLC cell lines included in the dependency map. (**C**) CRISPR gene effect of POU2F3, SMARCD1, BRD9, and SMARCA4 comparing POU2F3-positive SCLC lines vs. all other SCLC lines vs. all other cancer cell lines from the dependency map. 2-tailed unpaired t-test was used to generate p-values. n=4 POU2F3-expressing SCLC cell lines, n=19 other SCLC cell lines, n=1072 all other cancer cell lines. (**D-H**) Immunoblot analysis (D,E,G) and quantification of cell counts 3 and 6 days after plating (F,H) NCI-H1048 parental cells transduced with the indicated sgRNAs targeting SMARCD1, BRD9, BICRA/BICRAL, or non-targeting controls (sgControl). In D-F, POU2F3 sgRNAs are included as benchmark controls. For G, H, sgControl #1 was used. Cell counts are plotted as fold change relative to day 0. n=6 biological replicates for F and n=4 biological replicates for H. For all panels, *=p<0.05, **=p<0.01, ***=p<0.001 using a 2-tailed unpaired t-test. For all experiments, error bars represent mean +/− SEM.

Given the high degree of enrichment of mSWI/SNF and ncBAF components in our screen and the selective proliferative dependencies of POU2F3-positive SCLCs on ncBAF and mSWI/SNF components, we hereafter focused on these factors, especially given recently-achieved pharmacologic targeting of these remodelers. Multiple sgRNAs targeting both SMARCD1 and BRD9 significantly reduced POU2F3 protein levels and attenuated NCI-H1048 proliferation nearly to levels achieved by targeting POU2F3 itself (**Figure 2D-F**). Further, CRISPR-mediated suppression of GLTSCR1(*BICRA*) and GLTSCR1L(*BICRAL*), the nucleating paralog subunits of ncBAF complexes, similarly attenuated proliferation of the NCI-H1048 SCLC cell line (**Figure 2G,H**). Proliferative inhibition was further accentuated upon concomitant loss of both GLTSCR1/1L paralogs, which eliminates ncBAF complex assembly^41^. Together these data highlight that ncBAF complex integrity is required to maintain POU2F3 expression and proliferation in NCI-H1048 SCLC cells.

### Disruption of pan-mSWI/SNF ATPase and ncBAF-specific componentry attenuates SCLC proliferation

Given our findings above, we next asked whether small molecules that modulate mSWI/SNF complex activity or assembly impact the proliferation of POU2F3-positive SCLC. We employed two recently-discovered clinical-grade small molecules: FHD-609^42^, a VHL-based small molecule degrader of the BRD9 ncBAF complex subunit, and FHD-286^43^, a small molecule inhibitor of the SMARCA4/SMARCA2 mSWI/SNF ATPase subunits (**Figure 3A**). Notably, BRD9 degradation using FHD-609 and two additional BRD9 degrader compounds, WA-68-VQ71 and dBRD9A, reduced POU2F3 protein levels and attenuated proliferation of the two POU2F3-positive SCLC cell lines (NCI-H1048 and NCI-H211) with EC50 values in the low nanomolar range, but did not have an impact in the NCI-H526 and COR-L311 POU2F3-positive SCLC cell lines (**Figure 3B-D, S3A-C**). Further, treatment with FHD-286 and hence inhibition of mSWI/SNF ATPase activity resulted in dramatic reduction of POU2F3 protein levels and proliferative capacity with EC50 values in the low nanomolar range across all 4 POU2F3-positive SCLCs **(Figure 3E-G)**. These data were further validated using a potent SMARCA4/2 degrader, AU-15330^26^, and a second ATPase activity inhibitor, BRM014^36,44^ (**Figure 3E-G**). These phenotypes were selective for POU2F3-positive SCLC lines as SCLC cell lines of the ASCL1 and NEUROD1 subtypes were relatively insensitive to the BRD9 degraders, WA-68-VQ71 and FHD-609, and the SMARCA4/2 inhibitors FHD-286 and BRM014, with EC50 values >1 μM (**Figure S3D-F**). Treatment with these agents did not decrease ASCL1 or NEUROD1 protein expression in these cell lines except for modest changes in NEUROD1 in NCI-H82 cells and modest changes in ASCL1 in NCI-H1836 cells upon SMARCA4/2 inhibition treatment (**Figure S3G-J**). Together, these data underscore critical oncogenic maintenance functions of mSWI/SNF family complexes in POU2F3-expressing SCLC, supporting studies to mechanistically define these impacts and evaluate potential utility for these now clinical-grade agents. In addition, while all POU2F3-positive SCLCs are dependent on mSWI/SNF complex activity, these results indicate that further subdivision of POU2F3 SCLCs appears to demarcate sensitivity to ncBAF complex disruption.

**Fig. 3.**
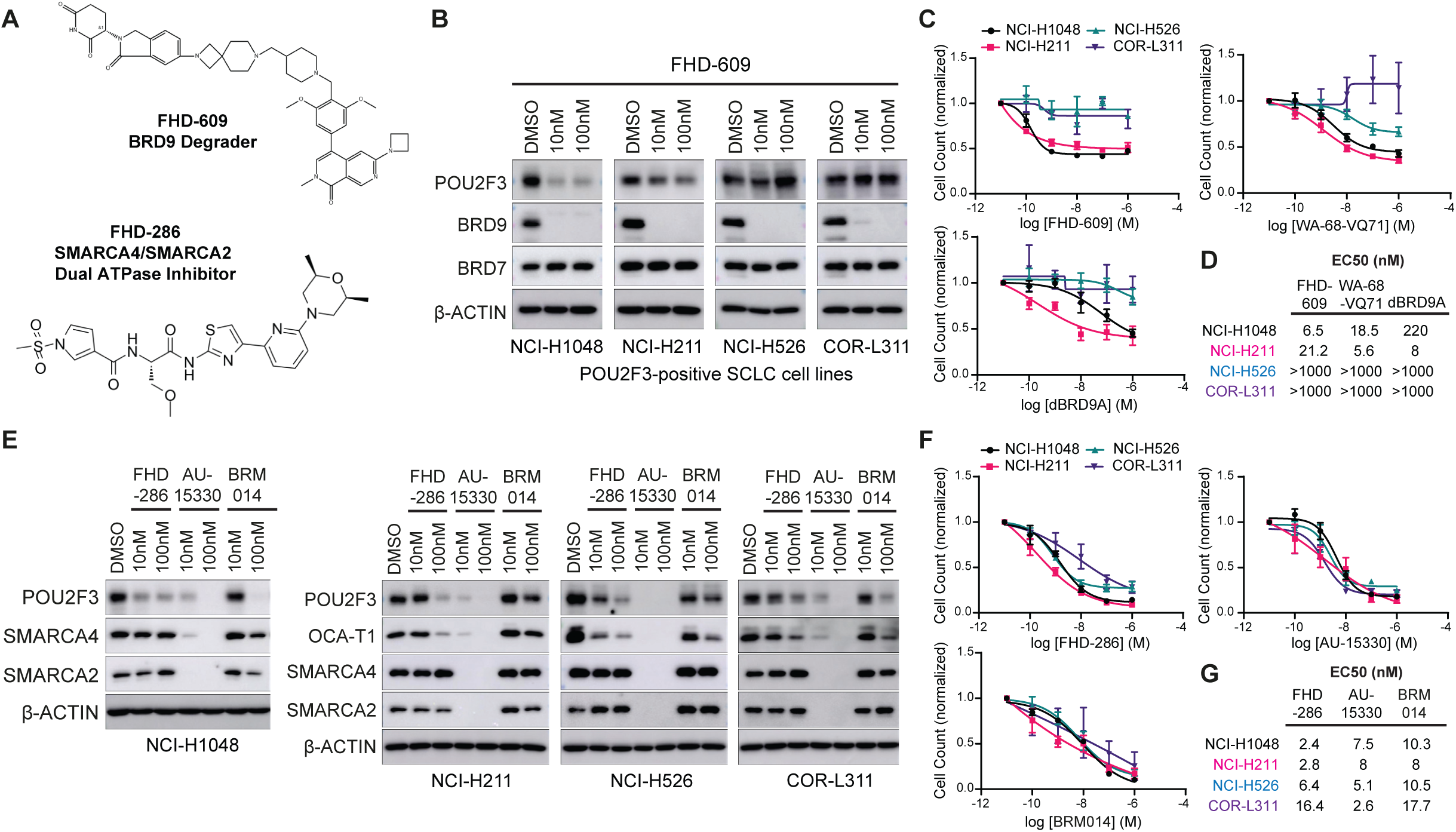
Disruption of pan-mSWI/SNF ATPase and ncBAF-specific componentry attenuates SCLC proliferation. (**A**) Structure of the clinical-grade BRD9 degrader FHD-609 and the clinical grade FHD-286 SMARCA2/4 ATPase inhibitor FHD-286. (**B**) Immunoblot analysis of the indicated POU2F3-expressing human SCLC cell lines after treatment with the BRD9 degrader (FHD-609) at the indicated concentrations or DMSO for 3 days. (**C**) Dose response assays on the indicated POU2F3-expressing human SCLC cell lines treated for 6 days with 3 different BRD9 degraders: FHD-609, WA-68-VQ71, or dBRD9A. (**D**) Table showing calculated EC50’s from the dose response curves from C. For C, n=3 biological replicates. (**E**) Immunoblot analysis of NCI-H1048, NCI-H211, NCI-526, and COR-L311 cells after treatment with SMARCA2/4 dual inhibitors (FHD-286 or BRM014) or the SMACRA2/4 degrader (AU-15330) at the indicated concentrations or DMSO for 3 days. (**F**) Dose response assays on the indicated POU2F3-expressing human SCLC cell lines treated for 6 days with FHD-286, AU-15330, or BRM014. (**G**) Table showing EC50’s from dose response assays from F. For F, n=3 biological replicates. For all doses response assays, cell counts are normalized to the DMSO condition. For all experiments, error bars represent mean +/− SEM. S.E.=short exposure and L.E.=long exposure.

### SCLC NE status tumors demarcate two subclasses and predict sensitivity to ncBAF disruption

We next sought to characterize the molecular differences between POU2F3-positive SCLC lines that do and do not exhibit sensitivity to ncBAF disruption. We performed RNA-seq experiments across the four cell lines, unsupervised clustering of which indicated clear segregation in the expression profiles of the NCI-H1048 and NCI-H211 cell lines and the NCI-H526 and COR-L311 cell lines (**Figure 4A**). Notably, genes corresponding to neuroendocrine features including *SYP* (synaptophysin)*, INSM1, CHGA* (chromogranin A)*, CHGB* (chromogranin B), *SEZ6, and ACTL6B* nearly exclusively demarcated the NCI-H526 and COR-L311 cell lines that were non-responsive to ncBAF inhibition, while those lines responding showed markedly reduced expression of these features (**Figure 4A**). Purely non-NE cell lines NCI-H1048 and NCI-H211 showed genes such as *LNMB1*, *CDCA3*, and *NKX6-1* uniquely upregulated and showed enrichment of immune signatures such as interferon alpha and gamma response, IL2 and IL6 JAK/STAT signaling, TGF-beta signaling and inflammatory response signaling pathways relative to the NE-like POU2F3-positive cell lines which were more enriched in oxidative phosphorylation, UV response and pancreas beta cell signatures (**Figure S4A).** Validation of selected NE-associated genes at the total protein level indicated clear separation between the purely non-NE and NE-like cell lines, with NE-like cell lines more closely mimicking those lacking POU2F3 expression (i.e. COR-L47) with respect to these features (**Figure 4B**).

**Figure 4.**
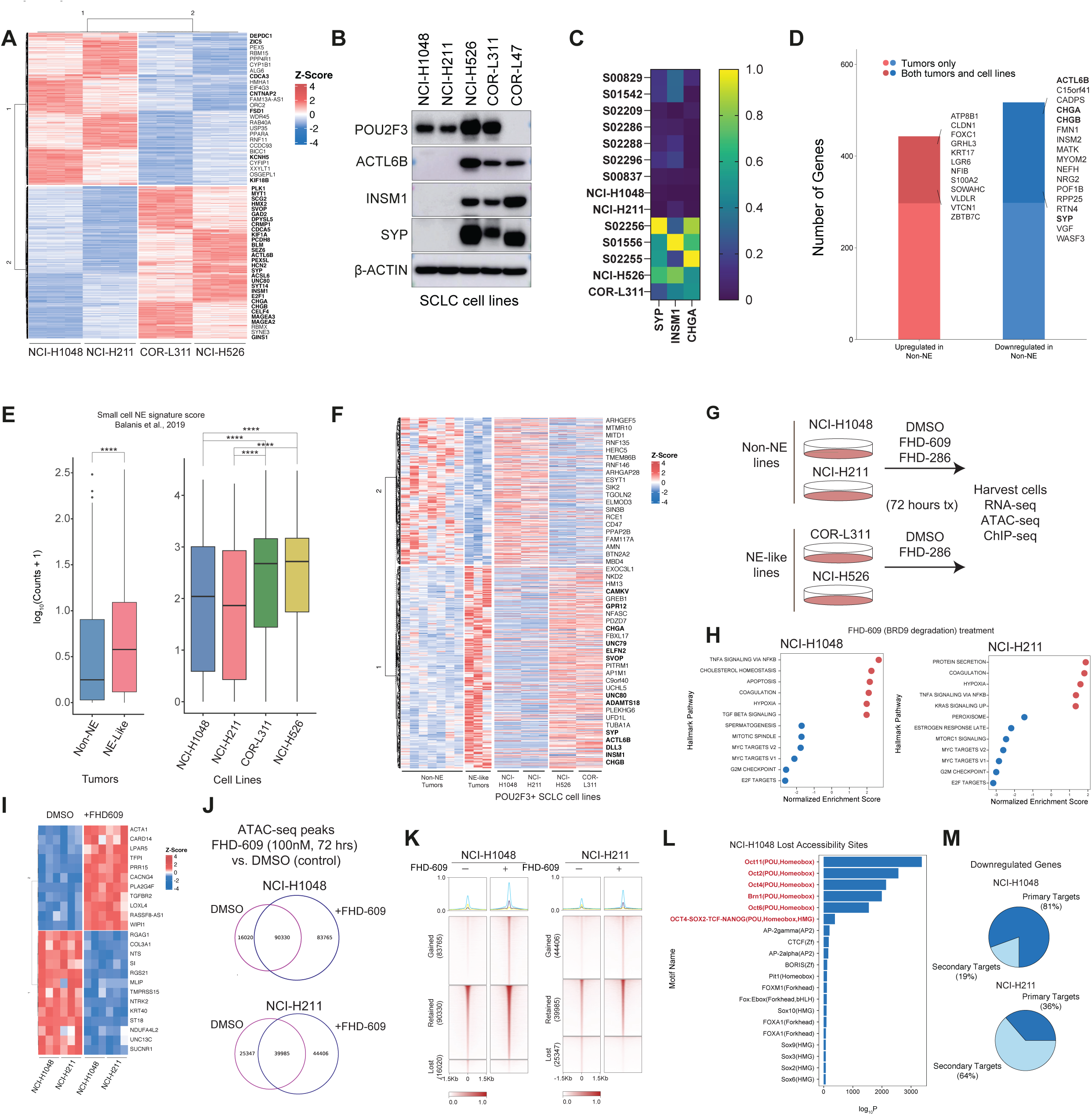
ncBAF disruption impacts non-NE-specific chromatin and gene regulation in POU2F3-positive SCLC. (**A**). Heatmap depicting top 10% upregulated and top 10% downregulated genes between NE-like and non-NE cell lines, by Z-score normalized expression. Bold indicates genes as being in the top 500 genes encompassed in Balanis et al., 2019 SCLC signature. (**B**) Immunoblot for ACTL6B, INSM1 and SYP NE markers performed on total protein isolated from SCLC cell lines. (**C**) Heatmap depicting gene expression of SYP, INSM1, and CHGA genes in SCLC primary tumors and cell lines. (**D**) Stacked bar graph depicting genes selectively up- and down-regulated in non-NE and NE-like SCLC primary tumors; dark red and dark blue indicate those genes overlapping genes selectively up- and down-regulated in SCLC cell lines. (**E**) Boxplot of log-transformed counts for top 500 SCNC signature genes in both George et al. 2015 mRNA data from SCLC tumors with POU2F3 expression and POU2F3-positive cell lines. Pairwise t-test was conducted to test for significance. * = p < 0.05 ** = p < 0.01 *** = p < 0.001 **** = p < 0.0001. (**F**) Heatmap (Z-score) showing top up-and down-regulated genes in the NE-like and non-NE primary tumors and cell lines. Genes bolded are those that overlap with the NE signature from Balanis et al., 2019. (**G**) Schematic depicting experimental strategy to evaluate mSWI/SNF modulatory small molecules in SCLC cell line models. (**H**) Dot plot depicting GSEA analysis Normalized Enrichment Scores (NES) for Hallmark pathways in FHD609- versus control-treated cells. (**I**) Heatmap depicting upregulated and downregulated genes shared between non-NE SCLC cell lines by differential gene analysis. Expression counts were z-score normalized by cell line. (**J**) Venn diagram depicting overlap between ATAC-seq peaks in FHD609-treated cells and control. (**K**) Heatmap showing gained, lost, and retained accessible peaks by ATAC-seq upon FHD-609 treatment in NCI-H1048 and NCI-H211 cells. (**L**) HOMER motif enrichment analysis performed on sites with lost accessibility upon FHD-609 treatment in NCI-H1048 cells. (**M**) Genes up- and down-regulated upon FHD-609 treatment that overlap with concordant changes in accessibility +/− 10kb (primary targets).

We next integrated these data with RNA-seq data from George et al. containing 12 primary SCLC tumors with POU2F3 expression^4^ (**Figure S4B**) to ask whether these two subsets exist within POU2F3-positive human SCLC, and to define gene signatures shared between primary tumors and our cell line models. Two POU2F3-positive tumors exhibited dominant expression of ASCL1 or NEUROD1 TFs relative to POU2F3 and thus were excluded for subsequent analyses (**Figure S4B, Table S2**). Of note, key NE marker genes such as *SYP*, *INSM1* and *CHGA* were expressed exclusively in the two NE-like cell lines and demonstrated much higher expression in a subset of n=3 POU2F3-positive primary tumors (**Figure 4C, S4C-E**). Indeed, PCA analyses confirmed several NE features, among others, driving this separation (**Figure S4E**). Gene expression analyses integrating both cell lines and primary tumor data indicated that among differentially expressed genes in primary SCLC tumors, 37% upregulated and 41% downregulated genes mirrored those selectively up or downregulated in our cell line models (**Figure 4D**). Genes such as *KDM5B*, *MAP3K5*, and *GRB7* were exclusively upregulated in the non-NE subgroup, while expression of genes such as *SYP*, *CHGA*, and *ACTL6B* were exclusively enhanced in the NE-like cell lines and primary tumors (**Figure 4D, S4D-E**). Further, using a neuroendocrine signature gene set of 250 total genes^45^, these two subgroups are even more clearly demarcated, indicative of NE features being the primary driver of the separation within the POU2F3-positive SCLC subtype (**Figure 4E-F, S4F**).

With these signatures and a potential molecular basis underpinning the diverse mSWI/SNF pharmacologic sensitivities, we next aimed to define the impact of either BRD9 degradation (FHD-609) or ATPase inhibition (FHD-286) on chromatin accessibility, histone landscape features, and subtype-specific gene expression (**Figure 4G**). To this end, we first subjected NCI-H1048 and NCI-H211 cell lines to treatment with either DMSO or FHD-609 (100nM) for 72 hours in culture. Gene expression (RNA-seq) analyses indicated top downregulated pathways in both cell lines included MYC target genes, G2M checkpoint, and E2F targets, in agreement with the strong antiproliferative impact we defined (**Figure 4H, S4G-I**, **Figures 2-3**). Overlap analyses performed between the two non-NE cell lines profiled defined several differentially expressed genes resulting from ncBAF disruption (FHD-609 treatment) that were common between the two cell lines, including *NTRK2*, *LPAR5*, and *TGFBR2* (**Figure 4I**).

To begin to define the underlying mechanisms, we next evaluated changes in chromatin accessibility using ATAC-seq. Treatment with FHD-609 for 72 hours revealed global changes, both decreases and increases, in accessibility genome-wide (**Figure 4J-K**). Notably, sites with reduction in chromatin accessibility upon FHD-609 treatment exhibited preferential enrichment for the POU2F3 DNA-binding motifs (POU homeobox motifs) (**Figure 4L, S4J**). Finally, we integrated these ATAC-seq and RNA-seq data to identify genes with nearby changes in accessibility (putative primary genes) and those without (putative secondary/downstream genes), finding that 81% and 36% of significantly downregulated genes were attributed to reduced nearby chromatin accessibility upon ncBAF disruption in NCI-H1048 and NCI-H211 cell lines, respectively (**Figure 4M**). Of note, among genes that we had defined as unique to the non-NE signature (**Figure 4A-B,E**) treatment with FHD-609 treatment more significantly led to changes in both accessibility and gene expression of those targets (**Figure S4K**). Taken together, these data highlight the impact of FHD-609-mediated ncBAF disruption at the chromatin accessibility and gene regulatory levels on the key pathways contributing to the maintenance of non-NE-like POU2F3-positive SCLC.

### mSWI/SNF ATPase inhibition attenuates distinct oncogenic gene regulatory networks mediated by distinct complexes

Given that all POU2F3-positive cell lines exhibited sensitivity to pan-mSWI/SNF ATPase inhibitors, we next sought to evaluate the impact of FHD-286 SMARCA4/2 inhibition across cell lines on gene and chromatin accessibility signatures as well as to define its effect on the genes that were impacted by ncBAF-disrupting small molecule degraders (FHD-609). In line with the antiproliferative impacts observed upon SMARCA4/2 ATPase inhibition (**Figure 3**), we identified strong downregulation of MYC target genes, E2F targets, and G2M cell cycle checkpoint across all four SCLC cell lines following FHD-286 treatment (**Figure 5A-B, S5A-D**). Overall impact on differential gene expression was larger relative to FHD-609 treatment in the non-NE SCLC cell lines, with over 1000 genes differentially down- and up-regulated upon treatment for 72 hours (**Figure 5C, S5A**). We identified largely separate groups of genes downregulated and upregulated by FHD-286 in non-NE and NE-like cell lines, however, genes such as *DUSP9*, *COL27A1*, and *MGAM2* were concordantly downregulated and *CARD14*, *MORN3* and *TGM2* others concordantly upregulated in both non-NE and NE-like cell lines (**Figure 5D**). ATAC-seq accessibility experiments revealed striking changes in the accessibility landscape across all 4 cell lines, most notably, losses in accessibility at over 30,000 sites genome-wide upon treatment with FHD-286 (**Figure 5E, S5E-F**). Decreases in accessibility were exemplified over the gene loci of tuft cell lineage markers *AVIL*, *CHAT*, *GFI1B*, and *SOX9* at which we also identified near complete loss of POU2F3 binding coupled with reduced occupancy of H3K27Ac and RNA Pol II^13,14^(**Figure 5F, S5G**). Notably, these lost sites were largely TSS-distal implicating altered enhancer accessibility and top motifs significantly enriched were those corresponding to the POU2F3 transcription factor, including POU homeobox factors (Oct1,2,6,11) (**Figure 5G, S5H)**. Integrating these accessibility findings with gene expression, we identified a collection of POU2F3 signature genes impacted via both chromatin accessibility and mRNA levels (**Figure 5H**). Taken together, these data show that mSWI/SNF ATPase inhibition uniformly impacts the accessibility over POU2F3 targets in POU2F3-positive SCLC, resulting in an attenuation of POU2F3 target chromatin accessibility and gene expression.

**Figure 5.**
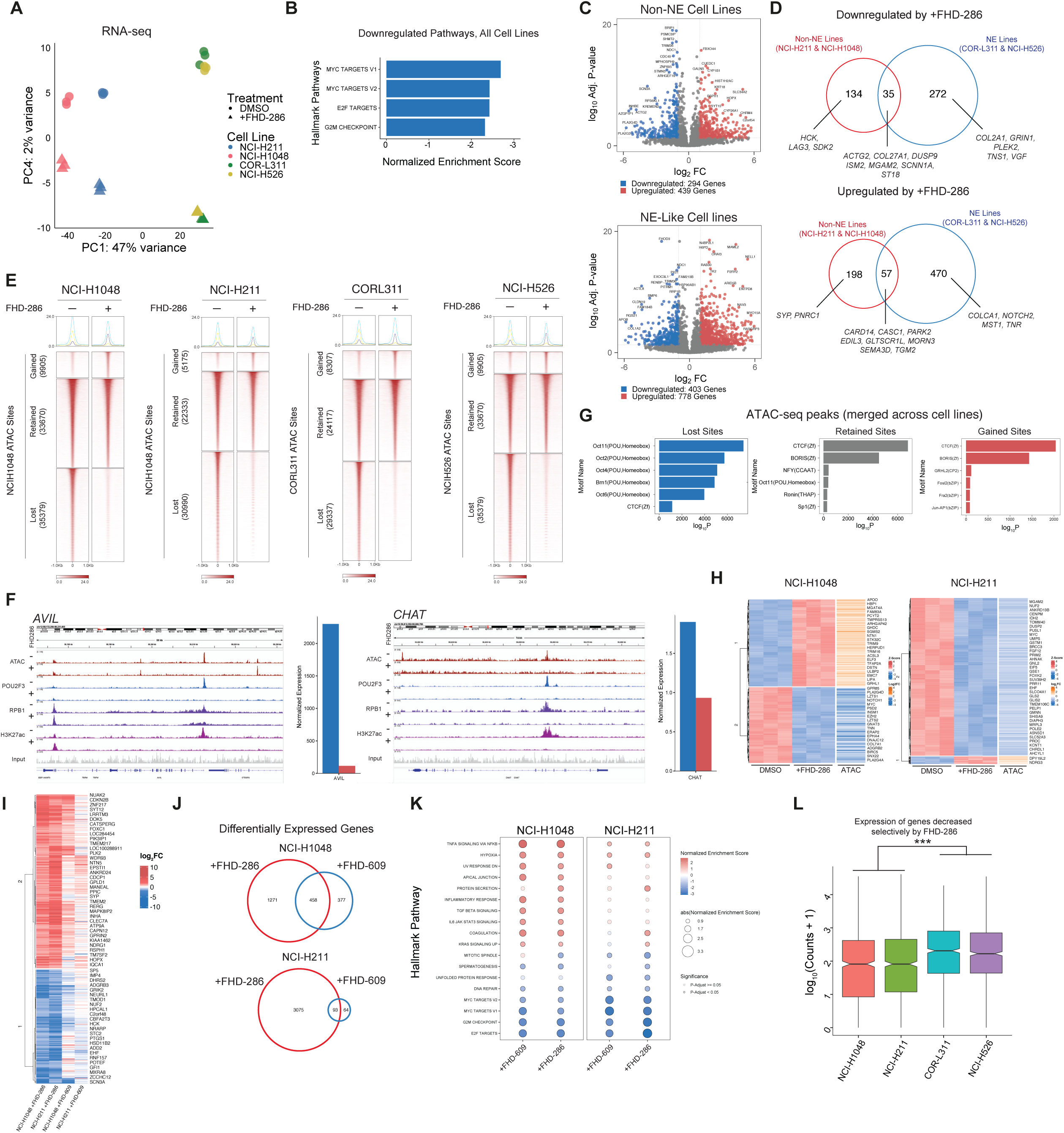
mSWI/SNF ATPase inhibition attenuates oncogenic SCLC gene regulatory networks. (**A**) PCA analysis performed on RNA-seq experiments in DMSO control and FHD-286-treated conditions across four POU2F3-positive SCLC cell lines. (**B**) GSEA analysis showing consistently and significantly downregulated pathways across all four SCLC cell lines. (**C**) Volcano plots showing significantly down- (blue) and up- (red) regulated genes in non-NE cell lines, NCI-H1048 and NCI-H211, and in NE-like cell lines, COR-L311 and NCI-H526 upon treatment with FHD-286. (**D**) Venn diagrams reflecting overlap between downregulated (top) and upregulated (bottom) genes impacted in non-NE and NE-like cell lines. Selected genes are labeled. (**E**) Heatmaps depicting lost, retained, and gained ATAC-seq peaks upon FHD-286 treatment called across the merged set of ATAC-seq peaks in each SCLC cell line. (**F**) Representative tracks at the AVIL and CHAT loci showing ATAC-seq signal, and POU2F3, RBP1, H3K27Ac ChIP-seq in DMSO and FHD-286 conditions. Bar graphs showing gene expression in DMSO and +FHD-286 conditions are shown to the right. (**G**) HOMER motif analyses performed over lost, retained, and gained ATAC-seq peaks across all cell lines. (**H**) Integrative RNA-seq and ATAC-seq analyses in DMSO- and FHD-286-treated conditions performed in NCI-H1048 and NCI-H211 cell lines shown as Z-scored (RNA) and LFC (ATAC) heatmaps. (**I**) Heatmap reflecting concordant changes in gene expression in the NCI-H1048 and NCI-H211 cell lines upon FHD-286 and FHD-609 treatments. (**J**) Venn diagrams depicting concordant differentially-expressed genes following FHD-286 and FHD-609 treatments. (**K**) Dot plot showing concordant and differential gene expression pathways impacted by FHD-286 and FHD-609 treatments. (**L**) Box and whisker plot depicting average expression of genes specifically impacted by FHD-286 treatment (FHD-286-only genes) in non-NE cell lines across all four POU2F3 SCLC cell lines.

We next sought to define the genes selectively impacted by FHD-286-mediated ATPase inhibition relative to FHD-609-mediated BRD9 degradation in the non-NE cell lines exhibiting sensitivity to both agents. We identified gene sets concordantly up- and down-regulated upon both FHD-609 and FHD-286 treatments in the NCI-H1048 and NCI-H211 cells, as well as genes and gene pathways that were only affected by FHD-286 pan-ATPase inhibition (**Figure S5I**). In general, a greater number of genes were impacted by SMARCA4/2 ATPase inhibition, and genes impacted by both small molecules were up or downregulated with higher amplitude upon FHD-286 relative to FHD-609 (**Figure 5I-J**). These included downregulated genes such as *NEURL1*, *CBFA2T3*, and *EHF* and upregulated genes such as *PLK2*, *DOK5*, and *SYT12* (**Figure 5I**). GSEA analyses revealed that concordantly downregulated genes included the expected MYC, E2F and G2M checkpoint targets while those specifically downregulated upon FHD-286 included mitotic spindle and glycolysis pathways (**Figure 5K**). Further, expression of genes that were specifically downregulated by FHD-286 treatment in the two non-NE cell lines was higher in the NE-like cell lines (COR-L311 and NCI-H526) that selectively responded to FHD-286 (and not FHD-609), demonstrating a gene level basis for the differential sensitivity we observed (**Figure 5L**). These data demonstrate the chromatin and gene regulatory impacts of FHD-286-mediated inhibition of mSWI/SNF complexes across POU2F3-positive SCLC and define sets of genes that are similarly and disparately regulated by ncBAF complexes and the mSWI/SNF family at-large.

### mSWI/SNF pharmacologic disruption impacts POU2F3 and co-activators OCA-T1/2

Treatment of all POU2F3-positive SCLC cell lines with SMARCA4/2 inhibitors (FHD-286, BRM014) and with BRD9 degraders (FHD-609, VA-68-VQ71) resulted in marked attenuation of expression of the POU2F3 gene signature defined by Vakoc and colleagues^13^ (**Figure 6A-B, S6A**). Of note, both treatments also dramatically attenuated signatures mediated by POU2F3 co-activators, OCA-T1 (*c11ORF53*) and OCA-T2 (*COLCA2*)^13^ (**Figure 6C-D, S6B**). Notably, FHD-286 treatment resulted in significantly reduced accessibility over the *POU2F3*, *c11orf53* and *COLCA2* gene loci across all cell lines (**Figure 6E, S6C,D**). Similarly, POU2F3 occupancy as well as occupancy of RNA Pol II and the H3K27Ac mark were reduced in signal at the *POU2F3* locus following FHD-286 treatment (**Figure 6E**). These data suggest a programmatic attenuation in the POU2F3-mediated gene signature that is achieved by pharmacologic inhibition of mSWI/SNF complexes.

**Figure 6.**
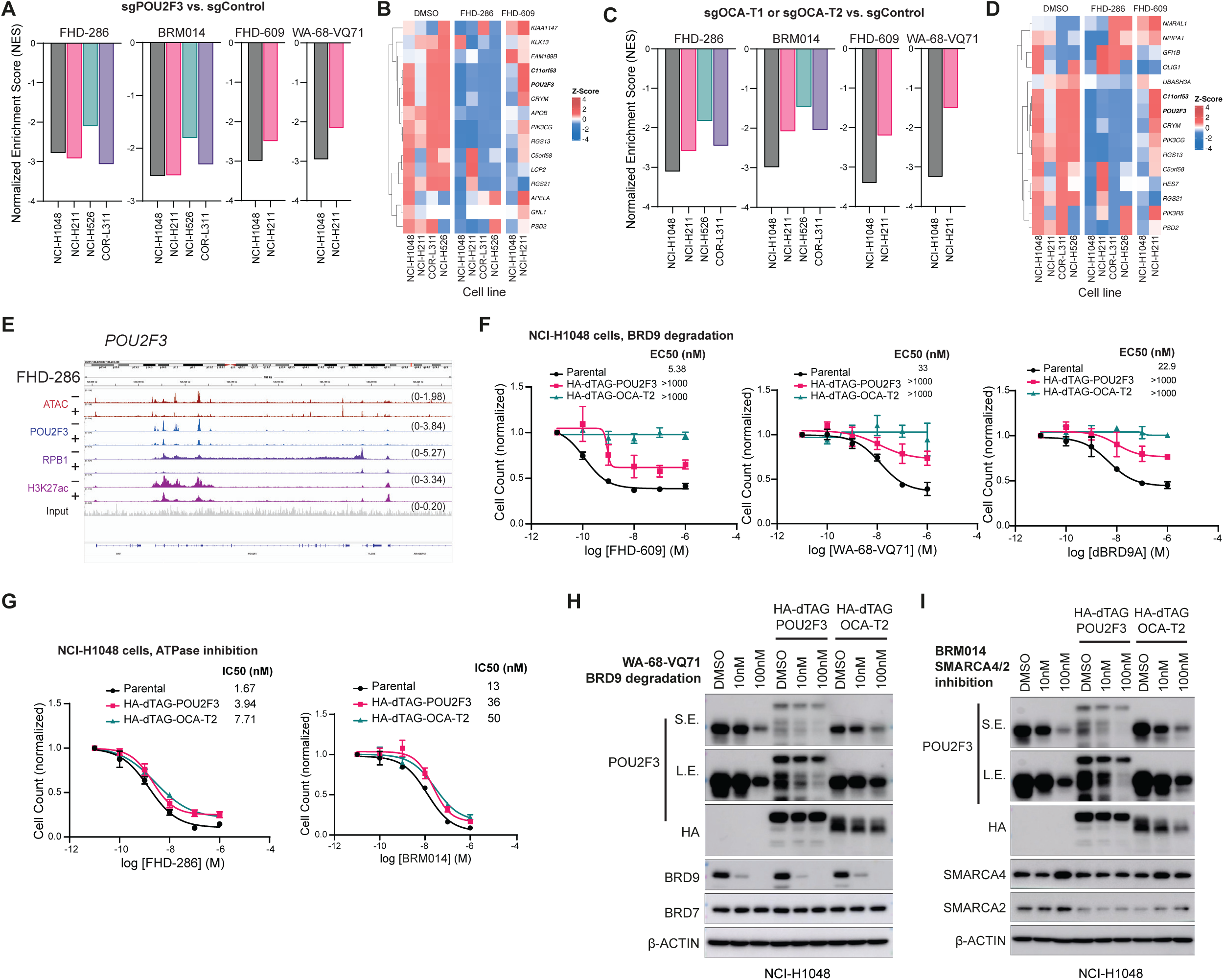
mSWI/SNF pharmacologic disruption impacts expression and activity of POU2F3 TF and co-activators OCA-T1/2. (**A,C**) Normalized Enrichment Score (NES) of downregulated genes after treatment with the SMARCA4/2 inhibitors (FHD-286 or BRM014) or BRD9 degraders (FHD-609 or WA-68-VQ71) at 100 nM for 72 hours comparing sgPOU2F3 vs. sgControl (A) or sgOCA-T1 (for NCI-H211, NCI-H526, COR-L311) or sgOCA-T2 (for NCI-H1048) vs. sgControl (C) from Wu et al., 2022. n=3 biological replicates for FHD-286 and FHD-609. n=2 biological replicates for BRM014 and WA-68-VQ71. (**B,D**) Heatmap depicting expression (Z-score (RPKM values) of sgPOU2F3 gene set signature genes (B) or sgOCA-T1 and sgOCA-T2 gene set signature genes (D) across all cell lines treated with either FHD-286 or FHD-609. (**E**) Chromatin targeting (ChIP-seq for POU2F3, RBP1, H3K27Ac) and accessibility regulation (ATAC-seq) at the POU2F3 locus in NCI-H1048 cells treated with FHD-286 (100 nM) for 72 hours. (**F**) Dose response assays of NCI-H1048 Cas9 cells expressing exogenous sgRNA-resistant HA-dTAG-POU2F3 with the knockout of endogenous POU2F3, or expressing sgRNA-resistant exogenous HA-dTAG-OCA-T2 with the knockout of endogenous OCA-T2 via CRISPR, or parental NCI-H1048 cells treated with FHD609 (left), the WA-68-VQ71 (middle), or dBRD9a (right) BRD9 degraders for 6 days. (**G**) Cells in F treated with FHD-286 (left) or BRM014 (right) for 6 days. For F,G, n=3 biological replicates; cell counts are normalized to the DMSO condition. Error bars represent mean +/− SEM. (**H-I**) Immunoblot analyses performed on cells from F,G treated with WA-68-VQ71 or BRM014 compounds at indicated concentrations for 72 hours.

Given these strong gene regulatory impacts of mSWI/SNF chemical disruption on putative TF drivers, we next sought to define the specific contributions of POU2F3 and co-activator (OCA-T2) expression to the maintenance of SCLC cell proliferation in culture. We used CRISPR/Cas9 to knockout endogenous POU2F3 or its co-activator OCA-T2^13^ and subsequently expressed exogenous, sgRNA-resistant POU2F3 or OCA-T2 under an EFS ubiquitous promoter fused to a degradation tag (dTAG)^46,47^ in NCI-H1048 cells (**Figure S6E-G**). We then performed 6-day dose response assays with three independent BRD9 degraders (FHD-609, WA-68-VQ71, dBRD9A) and two SMARCA4/2 ATPase inhibitors (FHD-286, BRM014) (**Figure 6F-G**). Strikingly, exogenous expression POU2F3 or OCA-T2 rendered SCLC cells uniformly resistant to all BRD9 degraders relative to parental NCI-H1048 cells, while not affecting efficacy of either SMARCA4/2 inhibitors (**Figure 6F-G**). Consistent with this, immunoblot analysis revealed that the BRD9 degrader WA-68-VQ71 decreased endogenous POU2F3, but did not alter exogenous POU2F3 or OCA-T2 protein levels (**Figure 6H**). In contrast, and again consistent with proliferation experiments, SMARCA4/2 inhibition (BRM014) decreased both endogenous POU2F3 and exogenous POU2F3 and OCA-T1 (**Figure 6I**). Together, these results demonstrate that loss of POU2F3 itself is responsible for the anti-proliferative effects observed with BRD9 degraders but that POU2F3 levels more modestly contribute to the anti-proliferative impact achieved with SMARCA2/4 inhibition, underscoring the added impact of pan-mSWI/SNF inhibition in POU2F3- positive SCLC.

### Clinical-grade pharmacologic disruption of mSWI/SNF complexes slows tumor growth and improves survival in POU2F3-positive SCLC

Lastly, we treated POU2F3-positive SCLC xenograft models (NCI-H1048 and NCI-H211) with either the BRD9 degrader, FHD-609, or the SMARCA4/2 inhibitor, FHD-286, both of which are currently in Phase I clinical trials. Pharmacodynamic (PD) experiments in NCI-H1048 xenografts demonstrated that FHD-609 completely degrades BRD9 protein with a corresponding decrease of its co-activator OCA-T2 (*COLCA2*) in all tumors (**Figure 7A-C**). FHD-286 also significantly decreased OCA-T2 expression relative to vehicle-treated mice (**Figure 7C**). We next performed efficacy studies in NCI-H1048 and NCI-H211 xenografts treated with FHD-609, FHD-286, or vehicle for 35 days continuously and then monitored off treatment for survival (**Figure 7D**). Importantly, we found that both FHD-609 and FHD-286 significantly inhibited tumor growth and increased survival relative to mice treated with vehicle in NCI-H1048 xenografts (**Figure 7E-F, S7A-B**). Similar results were observed in the NCI-H211 model (**Figure 7G-H, S7C-D**). Tumor growth and survival impacts occurred absent changes in body weight, suggesting that FHD-609 and FHD-286 were both well tolerated throughout the treatment course in both models (**Figure S7E-F**). Together, these *in vivo* results nominate clinical-grade mSWI/SNF-targeting small molecules as potential therapeutic agents in the treatment of the POU2F3 subtype of SCLC.

**Figure 7.**
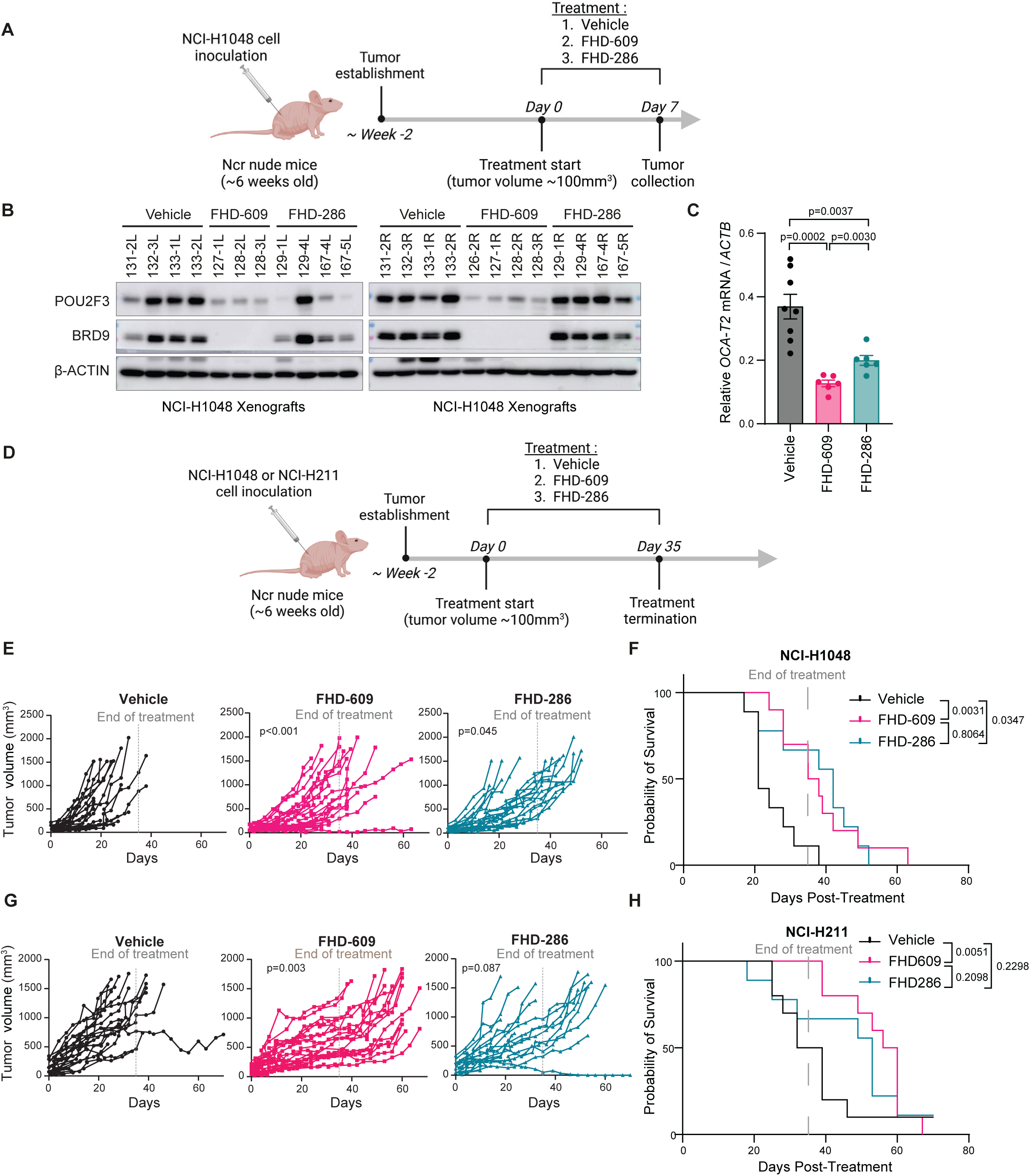
The BRD9 degrader FHD-609 or SMARCA4/2 inhibitor FHD-286 Slows Tumor Growth and Increases Survival in POU2F3 SCLC Xenograft Models. (**A**) Schema for the pharmacodynamic (PD) study in NCI-H1048 xenografts where Ncr nude mice harboring subcutaneous xenografts were treated for daily for 7 days with 0.5 mg/kg FHD-609 by intraperitoneal injection (IP), 1.5 mg/kg FHD-286 by oral gavage (PO), or vehicle (HP-β-CD). n=4 independent mice each with 1-2 tumors per mouse. (**B-C**) Immunoblot analysis (B) and RT-qPCR (C) of the tumors from A. For B, n=8 vehicle treated tumors, n=7 FHD-609 treated tumors, n=7 FHD-286 treated tumors. For C, n=8 vehicle treated tumors, n=6 FHD-609 treated tumors, and n=6 FHD-286 treated tumors. (**D**) Schema for the in vivo efficacy experiments in NCI-H1048 and NCI-H211 xenografts where Ncr nude mice harboring NCI-H1048 or NCI-H211 subcutaneous xenografts were treated continuously daily (QD) for 35 days with FHD-609 (0.5 mg/kg IP), FHD-286 (1.5 mg/kg PO), or vehicle (HP-β-CD), and then monitored off treatment for survival. (**E-F**) Spider plots of individual NCI-H1048 xenograft tumors (E) and Kaplan-Meier estimators of mice survival (F) treated with vehicle (Black), FHD-609 (0.5mg/kg IP QD) (pink) or FHD-286 (1.5mg/kg PO QD) (blue). For E,F, n=18 tumors from 9 independent mice (Vehicle), n=20 tumors from 10 independent mice (FHD-609), n=18 tumors from 9 independent mice (FHD-286). (**G-H**) Spider plots of individual NCI-H211 xenograft tumors (G) and Kaplan-Meier survival of mice (H) treated with vehicle (HP-β-CD) (Black), FHD-609 (0.5mg/kg IP QD) (pink) or FHD-286 (1.5mg/kg PO QD) (blue). For G,H; n=17 tumors from 10 independent mice (Vehicle), n=18 tumors from 10 independent mice (FHD-609), n=15 tumors from 9 independent mice (FHD-286). p-values are indicated on figure. For the spider plots in E,G, p-values are obtained from a linear mixed-effects model calculating the difference between the change in tumor volume relative to the vehicle arm (see Methods) while tumors were on treatment (until D35). For Kaplan-Meier survival curves in F and H, p-values are calculated using Gehan-Breslow-Wilcoxon test.

## Discussion

SCLC is a highly lethal form of lung cancer to date still lacking effective targeted therapies^2^. Studies over the past several years have demonstrated that SCLC maintenance relies on lineage-specific transcription factors including ASCL1, NEUROD1, and POU2F3, all of which currently remain undruggable targets^3,11–16,40^. Here, through a positive selection genome-wide CRISPR/Cas9 screen for POU2F3 regulators coupled with proliferation-based dependency screens performed across hundreds of cancer cell lines as well as validation studies^40^, we identified pivotal roles for mSWI/SNF chromatin remodeling complexes in the regulation of POU2F3 TF expression and maintenance of POU2F3-mediated oncogenic gene expression and proliferation. From this work, we find that all POU2F3-positive SCLCs are highly sensitive to the clinical-grade pan-mSWI/SNF inhibitor, FHD-286, including those that retain expression of canonical neuroendocrine markers. Surprisingly, we identify a subset of POU2F3-positive SCLCs that are purely non-neuroendocrine (non-NE lines) to be hypersensitive to ncBAF inhibition with clinical-grade and tool compound BRD9 degraders, including FHD-609. Together, our results suggest that pan-mSWI/SNF ATPase inhibition merits further evaluation as a therapeutic strategy for the broad class of POU2F3-positive SCLCs, while BRD9 degradation should be tested in POU2F3-positive SCLCs that are purely non-neuroendocrine as a potentially more selective strategy.

At the molecular level, we find that mSWI/SNF inhibition causes substantial loss of chromatin accessibility at POU2F3 target genes, including POU2F3 itself, implicating its autoregulation. Globally, ATPase inhibition via the FHD-286 SMARCA4/2 ATPase inhibitor results in a substantially more global ‘reset’ of the chromatin landscape, with the majority of affected sites reduced in accessibility, as expected, relative to FHD-609-mediated degradation of the ncBAF-specific subunit, BRD9. In non-NE SCLC cellular models, FHD-609-mediated ncBAF disruption led to an enhanced level of selectivity for accessibility losses over POU2F3 target sites, while pan-mSWI/SNF inhibition reduced accessibility over these sites as well as those corresponding to other TFs (**Figures 4,5**). These data highlight the SCLC addiction to the POU2F3 TF-driven signature enabled by mSWI/SNF-mediated chromatin accessibility generation; secondary gene pathways are supported broadly by mSWI/SNF family chromatin remodelers, rather than the ncBAF subcomplex in isolation. These findings are further supported by rescue experiments involving exogenous POU2F3 or OCA-T2 expression that we show selectively rescues the cellular proliferation defects caused by BRD9 degraders but not those caused by pan mSWI/SNF complex ATPase inhibition.

Human POU2F3-positive SCLC has historically been considered to be a purely non-neuroendocrine subtype of SCLC^3,7^. Here we found that all POU2F3-positive human SCLC cell lines were selectively hypersensitive to inhibition of mSWI/SNF, but only some cell lines were hypersensitive to disruption of ncBAF through BRD9 degradation. Unbiased analyses to identify transcriptional signatures associated with BRD9 dependence in POU2F3-positive cell lines uncovered gene signatures associated with ncBAF sensitivity, which included differences in neuroendocrine marker expression. Specifically, POU2F3-positive cell lines that retained expression of neuroendocrine markers were inherently resistant to BRD9 degradation, while cell lines with complete absence of neuroendocrine markers were hypersensitive to BRD9 degradation. Similar to our observations in cell lines, RNA-seq data from human POU2F3-positive SCLC patients^4^ also showed that POU2F3-positive human SCLC segregates into mostly pure non-neuroendocrine tumors with fewer POU2F3-positive tumors retaining some expression of neuroendocrine markers. Our findings suggest potential therapeutic relevance for further subclassification of the POU2F3 subtype into pure non-neuroendocrine tumors or tumors that retain neuroendocrine marker expression, especially relevant given that BRD9 degraders and SMARCA2/4 inhibitors are currently being tested in clinical trials for other cancer indications. Deciding on the appropriate therapeutic approach in non-neuroendocrine POU2F3-positive SCLCs will require additional efforts to evaluate toxicities of each therapeutic strategy to maximize therapeutic window. Future studies will be needed to address whether there are biological differences and unique therapeutic opportunities correlated with expression or absence of neuroendocrine markers within the POU2F3 subtype. Moreover, given the limited number of POU2F3-positive human SCLCs available with transcriptome sequencing, future studies will also require a larger number of samples to further refine mechanistic findings in the setting of transcriptional heterogeneity.

Our data show anti-tumor efficacy of BRD9 degraders or SMARCA4/2 inhibitors in POU2F3-positive SCLC xenograft models without overt toxicity. POU2F3-positive SCLC is the least responsive SCLC subtype to first-line chemotherapy^8^ suggesting that new upfront therapeutic approaches for the POU2F3-subtype are needed either in lieu of chemotherapy or in combination with chemotherapy. We observed meaningful anti-tumor activity with BRD9 degraders and with SMARCA4/2 inhibitors as monotherapy, but we did not observe striking tumor regressions suggesting that combination strategies should be considered with chemotherapy or with other targeted agents that may synergize with BRD9 or SMARCA4/2 without overlapping toxicity. Interestingly, ncBAF complexes have been implicated in homologous recombination and DNA repair^48^. BRD9 inactivation has been shown to sensitize ovarian cancer cells to PARP inhibition^48^, which is particularly relevant given that POU2F3-positive SCLCs are highly sensitive to PARP inhibitors^8^. The identification of effective combination strategies that enhance activity of mSWI/SNF inhibition or BRD9 degradation in POU2F3-positive SCLC represent important next investigations.

Finally, recent evidence is emerging that SCLC subtypes could be regulated by distinct epigenetic modifiers. Our data suggest that only the POU2F3 subtype is highly dependent on mSWI/SNF; in parallel, previous studies found that some SCLC cell lines of the ASCL1 subtype are highly dependent on the histone demethylase LSD1, where LSD1 is dispensable in other SCLC subtypes including in POU2F3-positive SCLC^49,50^. As such, the identification of the biochemical and chromatin regulatory interplay between lineage-specific TFs and each distinct chromatin regulatory entity is likely to yield improved understanding regarding SCLC pathogenesis and uncover additional therapeutic vulnerabilities associated with each SCLC subtype.

### Limitations of Our Study

While we identify and validate that mSWI/SNF complexes represent top-ranked, chemically targetable vulnerabilities in POU2F3-positive SCLC, the number of available cell lines and *in vivo* models for this cancer remains limited. Further, the programmatic changes in oncogenic gene expression upon mSWI/SNF disruption described here very likely exhibit increased variability in human tumors, presenting challenges in the comprehensive identification of reliable ‘biomarkers’ that faithfully predict either sensitivity and response to mSWI/SNF ATPase inhibitors or BRD9 degradation. Further, given that these SCLC cell lines have not been subjected to whole-genome sequencing, it remains unclear whether additional non-coding mutations in cis-regulatory elements contribute alone or in combination with mSWI/SNF complexes to regulate POU2F3 expression in SCLC. Finally, in our unbiased CRISPR/Cas9 screen using a positive selection assay in NCI-H1048 cells, we identified several genes that when inactivated decreased POU2F3 expression, which we validated in low-throughput knockout experiments. However, we cannot rule out the possibility of false negatives given that this was a positive selection assay in a cell line with a POU2F3 dependency.

## Acknowledgements

We thank members of the Oser and Kadoch labs for their critical feedback. We are grateful to members of the DFCI Molecular Biology Core Facility, including Zach Herbert and Maura Berkeley. We also thank members of the Barbie and Jänne labs for helpful discussions. This work was supported a Novartis DDTRP grant (M.G.O.), NCI R37CA269990 (M.G.O.), and a William Raveis Charitable Fund Damon Runyon Clinical Investigator Award (M.G.O) supported by the Damon Runyon Cancer Research Foundation (CI-101-19; M.G.O), the Mark Foundation for Cancer Research Emerging Leader Award (C.K.) and the Howard Hughes Medical Institute (C.K.). K.S. was supported by the National Science Foundation Graduate Research Fellowship Program.

## Author Contributions

L.D. and K.S.: Conceptualization, methodology, validation, formal analysis, investigation, data curation, writing – original draft, visualization, project administration. X.L.: investigation, methodology, validation, formal analysis, data curation. A.W.Y., Y.L., X.Q., R.L., S.S..: Formal analysis, data curation, visualization, methodology. X.S.W: methodology, investigation. Q.L., J.Q.: chemical synthesis and resources. C.R.V, H.W.L.: resources. H.H., E.M: formal analysis. Y.L., J.G.D: methodology. T.D.D.S, W.C.F, T.Z., T.A., M.J.N.: resources. C.R.V., H.W.L.: supervision, resources. C.K. and M.G.O: Conceptualization, methodology, investigation, resources, data curation, writing – original draft, visualization, supervision, project administration, funding acquisition.

## Inclusion and Diversity Statement

We support inclusive, diverse and equitable conduct of research. One or more of the authors of this paper self-identifies as an underrepresented ethnic minority in their field of research or within their geographical location.

## Methods

### SCLC Cell Lines

NCI-H1048, NCI-H526 (09/2019), NCI-H211 (01/2022), COR-L311 (01/2023), NCI-H1092, NCI-H1836, LU135 (07/2022), and 293T cells were obtained from American Type Culture Collection (ATCC). SCLC-22H were purchased from DSMZ (01/2023). COR-L47 were obtained from Sigma (11/2018). NCI-H82 cells were a kind gift from Dr. Kwok-kin Wong’s laboratory (New York University) and were obtained in 8/2014. NCI-H526, NCI-H211, COR-L311, COR-L47, NCI-H82 and LU135 cells were maintained in RPMI-1640 media with 10% FBS and P/S. NCI-H1048 cells were maintained in RPMI-1640 media with 10% FBS, P/S and ITS. NCI-H1092 and NCI-H1836 cells were maintained in DMEM/F12 media 5% FBS, P/S, and HITES. SCLC-22H and 293T was maintained in DMEM media with 10% FBS and P/S. Early passage cell lines were tested for Mycoplasma (Lonza #LT07-218) and then were frozen using Bambanker’s freezing media (Bulldog Bio). All experiments were performed with cell lines that were maintained in culture for <4 months at which time an early passage cell line was thawed.

### Homology-directed repair using the Alt-R™ CRISPR-Cas9 System and HDR Donor

#### Double-stranded HDR DNA Template Production

The DCK*-P2A-GFP sequence was obtained performing two distinct PCR reactions using KOD HOT START DNA POLYMERASE (Fisher Scientific #710863) and the pLX304-DCK*-IRES-GFP^37^ vector as a template for overhang PCR that introduced linker1 and P2A onto the 5’ and 3’ ends of DCK* or P2A and linker2 onto the 5’ and 3’ ends of GFP to obtain 2 PCR products: 1. linker1-DCK*-P2A and 2. P2A-GFP-linker2 sequence. DCK* is a variant of the deoxycytidine kinase with Ser74Glu, Arg104Met, and Asp133Ala substitutions. Then, an overlap PCR reaction was performed to obtain the linker1-DCK*-P2A-GFP-linker2 (called DCK*-P2A-GFP) sequence. In parallel, 2 double strand DNA sequences coding for the homology arms of *POU2F3* were designed and ordered from IDT as gBlocks™ Gene Fragments. Homology arm 1 is 300 nucleotides upstream of the endogenous stop codon of human *POU2F3* gene flanked by M13 pUC and Linker1 sequence (M13-HA1-linker1 called HA1). Homology arm 2 is 300 nucleotides downstream of the human *POU2F3* gene flanked by linker2 and M13 reverse sequence (Linker2-HA2-M13Rv called HA2). gBlocks™ (500 ng) were spin down before being resuspended at ∼50 ng/µL in IDTE Buffer and incubated at 50°C for 15-20 minutes. An in-fusion cloning reaction was then performed using all 3 fragments above (HA1, HA2, and DCK*-P2A-GFP) mixed with a purified inverse PCR product of the pDONR223 backbone that was made using M13 forward and M13 reverse primers. For the In-Fusion reaction, a 1:2 plasmid:insert (DCK*-P2A-GFP/HA1/HA2) ratio was used [121.8 fmol (200ng) of the pDONR223 PCR product mixed with 243 fmol of each dsDNA insert sequences] and then mixed with 2µl of In-Fusion 5X HD Cloning Plus (Takara Bio #638909). The reaction was incubated for 15 minutes at 37°C and stored at −20°C. The reaction mixture was then transformed at a ratio of 1:10 (reaction volume/volume competent cells) into HB101 competent cells (Promega #L2011). Spectinomycin-resistant colonies were screened by restriction digestion of miniprep DNA and subsequently validated by gel electrophoresis in a 0.8% agarose gel and by DNA sequencing. Finally, PCR reaction was performed on the new synthetized vector to amplify the HA1-DCK*-P2A-GFP-HA2 sequence corresponding to the HDR (homology directed repair) DNA template and was purified using the NucleoSpin PCR Clean-up kit (Macherey-Nagel # 740609.10) according to the manufacturer’s protocol. All DNA sequences used in the study are listed in Supplementary Table 3.

#### RNP production and nucleofection

The RNP complex was produced by complexing a two-component gRNA to Cas9 according to the homology-directed repair (HDR) protocol from IDT. Briefly, crRNA was designed using both the Broad Institute sgRNA designer tool (http://portals.broadinstitute.org/gpp/public/analysis-tools/sgrna-design) and the IDT sgRNA designer tool (https://www.idtdna.com/site/order/designtool/index/CRISPR_SEQUENCE) predicted to cut at a maximum of 6 nucleotides from the insertion site (before the endogenous stop codon of POU2F3 gene) and synthesized by IDT technologies. The following crRNA oligo predicted to cut 1 nucleotide before the insertion site was used: 5’-AAACTTTTTGGTCTCAGTGG-3’ (antisense). To prepare gRNA complex, the designed crRNA and a TracrRNA (IDT# 1072533) were resuspend at 100µM in Nuclease-free duplex buffer, mixed 1:1 by volume and annealed by incubation at 95°C for 5 min. Then, RNP complex was prepared by mixing gRNA complex with Alt-R® S.p. Cas9 Nuclease V3 (IDT #1081059) at a ratio 2:3 by volume and incubated 20 min at room temperature. RNP was electroporated immediately after complexing. In parallel, 1x10^6^ NCI-H1048 cells were prepared, washed in sterile PBS, and resuspended in 20 µL of SF Cell Line Nucleofector™ Solution with the supplement added (Lonza V4XC-2032). The cells and the RNP complex were carefully mixed and transferred to a well of the 16-well Nucleocuvette™. Nucleofection was performed using CM-137 program on 4D-Nucleofector™ (Lonza). Cells were carefully resuspended with pre-warmed media, transferred into a 12 well plate and incubated at 37°C, 5% CO_2_. Cells were amplified for several days to allow FACS sorting for GFP-positive cells.

#### Validation of NCI-H1048 POU2F3-DCK*-P2A-GFP Knock-in cells

NCI-H1048 cells nucleofected with the POU2F3-DCK*-P2A-GFP dsDNA template were sorted twice by FACS for GFP expression to obtain a pure population expressing POU2F3-DCK*-P2A-GFP fusion protein. Endogenous expression of POU2F3-DCK* fusion protein and loss of the WT POU2F3 protein was confirmed by immunoblot analysis for POU2F3. Specificity of the knock-in for the *POU2F3* locus was verified by blotting for DCK as the DCK immunoblot only showed 1 additional band, apart from endogenous DCK, at the expected molecular weight of the POU2F3-DCK*-P2A-GFP fusion with an identical molecular weight also seen on the POU2F3 immunoblot. Functionality of POU2F3-DCK* fusion was verified by BVdU dose response assays and rescue experiments with sgRNAs targeting DCK.

### Flow Cytometry

NCI-H1048 parental and NCI-H1048 POU2F3-DCK*-P2A-GFP cells were collected, washed twice in PBS, resuspended in FACS buffer (D-PBS containing 2% FBS) and, transferred to flow cytometry tubes containing a 70 μm filter and analyzed for GFP expression on a LSR Fortessa flow cytometer (Becton Dickinson, Franklin Lakes, NJ). Data were analyzed using FlowJo software.

### BVdU Sensitivity and Rescue Experiments

NCI-H1048 parental and NCI-H1048 POU2F3-DCK*-P2A-GFP cells were seeded into twelve-well plates at 50,000 cells per well in 0.5 mLs of complete media. Each well in the twelve-well dish received 0.5 ml of a stock solution of BVdU to achieve final BVdU concentrations of 1 µM, 10 µM, 100 µM, 200 µM and 500 µM. A total of 0.5 mLs of media with DMSO was added to the sixth well as a control. Seven days later, the cells were collected and counted using a Vi-Cell XR cell counter. The same protocol was used for DCK sgRNA rescue experiments using NCI-H1048 parental and NCI-H1048 POU2F3-DCK*-P2A-GFP cells infected with a sgRNA targeting DCK or a non-targeting sgRNA and treated with BVdU (10 µM) or DMSO for 7 days.

### BVdU-Positive Selection CRISPR/Cas9 Screen

On day 0, NCI-H1048 POU2F3-DCK*-P2A-GFP cells were counted. 1.2 x 10^8^ cells (which would yield representation of ∼500 cells/sgRNA) were pelleted and resuspended at 2 x 10^6^ cells/mL in complete media containing 50 µl/mL of the Cas9-containing whole genome Brunello sgRNA library (CP0043)^38^ (Purchased from the Broad Institute) lentivirus and 8 µg/mL polybrene. The lentiviral titer was determined empirically in pilot experiments with a goal multiplicity of infection (MOI) of 0.3-0.5. CP0043 contains 77,441 sgRNAs targeting 4 sgRNAs per gene with 1000 non-targeting sgRNAs controls. The cells mixed with polybrene and lentivirus were then plated in 1 mL aliquots onto 12 well plates and centrifuged at 2000 rpm (931 x g) [Allegra® X-15R Centrifuge (Beckman Coulter), rotor SX4750A] for 2 hours at 30°C. In parallel, 2 x 10^6^ cells were also spin infected under the same conditions but without lentivirus as a control for puromycin selection (Mock). ∼16 hours later (day 1), the cells were collected, pooled, and centrifuged to remove the lentivirus and polybrene, and the cell pellet was resuspended in complete media at 0.5 x 10^6^ cells/mL and plated into ten 15 cm tissue culture treated plates at 0.34 x 10^6^ cells/mL or in 6 well plates for the control (Mock) cells. The cells were then cultured for 48 hours at which time (day 3) the cells were counted and plated at 0.4 x 10^6^ cells/mL in 15 cm tissue culture treated plates with fresh media containing puromycin (0.25 µg/mL) to select for puromycin-transduced cells. A parallel experiment was performed on day 3 to determine the MOI of the screen. To do this, the cells infected with the sgRNA library or mock-infected cells were plated at 0.4 X 10^6^ cells/mL in 6 well plates in the presence or absence of puromycin (0.25 µg/mL). After 72 hours (day 6), cells were counted using the Vi-Cell XR Cell Counter and the MOI was calculated using the following equation: (# of puromycin-resistant cells infected with the sgRNA library/# total cells surviving without puromycin after infection with the sgRNA library) – (# of puromycin-resistant mock-infected cells/ # total mock-infected cells). The actual MOI was 0.56 for biological replicate 1 and 0.52 for biological replicate 2. On day 7, all puromycin-resistant cells were pooled, collected and counted. A total of 2 x 10^8^ cells were maintained to pursue the screen and were plated again at 0.4 x 10^6^ cells/mL with fresh media. On day 9, cells were pooled, counted and a total of 4 x 10^7^ cells for each condition (therefore maintaining at least 500 cells/guide) and were plated at 0.05 x 10^6^ cells/mL in 37 15cm tissue culture plates with complete media containing BVdU (10 µM) or at 0.3 x 10^6^ cells/mL in 6 15cm tissue culture plates with complete media for the untreated arm. At the same time, the remaining cells were collected and divided in aliquots of 4 x 10^7^ (again to maintain representation of at least 500 cells/guide), washed in PBS, and cell pellets were frozen for genomic DNA isolation for the initial timepoint prior drug selection. On days 12 and 14, the cells from the untreated arm were pooled, counted, and a total of 4 x 10^7^ cells for each replicate were plated back at 0.3 x 10^6^ cells/mL into 6 15cm tissue culture plates with fresh complete media. On day 16, 7 days after BVdU treatment, the screen was ended by collecting all remaining cells in both the BVdU and untreated conditions. As above, the remaining cells were divided in aliquots of 4 x 10^7^ cells, washed in PBS and cells pellets were frozen for genomic DNA isolation. The screen was performed in two biological replicates each with a separate infection.

Following completion of the screen, genomic DNA was isolated using the Qiagen Genomic DNA maxi prep kit (cat. # 51194) according to the manufacturer’s protocol. Raw Illumina reads were normalized between samples using: Log2[(sgRNA reads/total reads for sample) × 1e6) + 1]. The initial common time point data (day 9) was then subtracted from the end time point after BVdU selection or in the untreated arm (day 16) to determine the relative enrichment of each individual sgRNAs in the BVdU arm vs. untreated arm over time. The subtracted Log2 normalized reads of the BVdU arm (day 16) vs. the untreated arm (day 16) were then analyzed using Hypergeometric and STARS analysis comparing the average of the BVdU treated samples at day 16 vs. the average of the untreated samples at day 16. Analysis of the individual replicates were also performed each yielding similar results to the average of both biological replicates. Apron analysis was also performed using the normalized counts of replicates 1 and 2 of the BVdU treated samples at day 16 compared to the average of the untreated samples at day 16. Apron, Hypergeometric, and the STARS analyses were done using the GPP web portal (https://portals.broadinstitute.org/gpp/screener/). The averaged data from 2 biological replicates were used for all analyses shown in the manuscript.

### sgRNA Cloning to Make Lentiviruses

sgRNA sequences were designed using the Broad Institute sgRNA designer tool (http://portals.broadinstitute.org/gpp/public/analysis-tools/sgrna-design) for the DCK sgRNA rescue experiment or sgRNA sequences were used from the Brunello sgRNA library. All oligos were synthesized by IDT technologies. The sense and antisense oligonucleotides were mixed at equimolar ratios (0.25 nanomoles of each sense and antisense oligonucleotide) and annealed by heating to 100°C in annealing buffer (1X annealing buffer 100 mM NaCl, 10 mM Tris-HCl, pH 7.4) followed by slow cooling to 30°C for 3 hours. The annealed oligonucleotides were then diluted at 1:400 in 0.5X annealing buffer.

For CRISPR/Cas9 knockout experiments in cells, the annealed oligos were ligated into LentiCRISPRV2-Puro (Addgene # 98290) or lentiCRISPRV2-Neo (Addgene #98292 where the puromycin resistance gene was replaced with the G418 resistance gene) for experiments where double knockout is needed. Ligations were performed with T4 DNA ligase for 2 hours at 25°C. The ligation mixture was transformed into HB101 competent cells. Ampicillin-resistant colonies were screened by restriction digestion of miniprep DNAs and subsequently validated by DNA sequencing.

The following sgRNA oligos (including the BsmBI sites) were used to clone into the LentiCRISPR V2-Puro vector for CRISPR knockout experiments: *DCK* human sense (5’-CACCGTCGAAGGGAACATCGCTGCA-3’), *DCK* human anti-sense (5’-AAACTGCAGCGATGTTCCCTTCGAC-3’), *POU2F3* human sense #1 (5’-CACCGTCCTACCAAATACTTCACTG-3’), *POU2F3* human anti-sense #1 (5’-AAACCAGTGAAGTATTTGGTAGGAC-3’), *POU2F3* human sense #2 (5’-CACCGATCACAGTGTTACCTGACAT-3’), *POU2F3* human anti-sense #2 (5’-AAACATGTCAGGTAACACTGTGATC-3’), *SMARCD1* human sense #1 (5’-CACCGGAAACGGCTAGATATCCAAG-3’), *SMARCD1* human anti-sense #1 (5’-AAACCTTGGATATCTAGCCGTTTCC-3’), *SMARCD1* human sense #2 (5’-CACCGTTTGTCCAGTTCAATCACCA-3’), *SMARCD1* human AAACTGGTGATTGAACTGGACAAAC-3’), *SMARCD1* human CACCGTGACAAACTCCCGCTCGTGA-3’), *SMARCD1* human AAACTCACGAGCGGGAGTTTGTCAC-3’), *SMARCD1* human CACCGGGCAGCCGAATGACACCTCA-3’), *SMARCD1* human AAACTGAGGTGTCATTCGGCTGCCC-3’), *SMARCD1* human CACCGGGGAGCTTCGGGTAGAAGGA-3’), *SMARCD1* human AAACTCCTTCTACCCGAAGCTCCCC-3’), *SMARCD1* human CACCGGACCTGCTGGATCTGCTGAG-3’), *SMARCD1* human AAACCTCAGCAGATCCAGCAGGTCC-3’), *EP300*human sense #1 (5’-CACCGATGGTGAACCATAAGGATTG-3’), *EP300*human anti-sense #1 (5’-AAACCAATCCTTATGGTTCACCATC-3’), *EP300*human sense #2 (5’-CACCGGTGGCACGAAGATATTACTC-3’), *EP300*human anti-sense #2 (5’-AAACGAGTAATATCTTCGTGCCACC-3’), *MED19*human sense #1 (5’-CACCGTCCTGGTCCGAAGCCGAGTG-3’), *MED19*human anti-sense #1 (5’-AAACCACTCGGCTTCGGACCAGGAC-3’), *MED19*human sense #2 (5’-CACCGCACCTGGCAGTTCCCTCATG-3’), *MED19*human anti-sense #2 (5’-AAACCATGAGGGAACTGCCAGGTGC-3’), *IPPK* human sense #1 (5’-CACCGGTCCCCTTGATCTCTACTCA-3’), *IPPK* human AAACTGAGTAGAGATCAAGGGGACC-3’), *IPPK*human CACCGAAGGACCTGGATACTCTCAG-3’), *IPPK* human AAACCTGAGAGTATCCAGGTCCTTC-3’), *KAT7* human CACCGATGAACGAGTCTGCCGAAGA-3’), *KAT7* human AAACTCTTCGGCAGACTCGTTCATC-3’), *KAT7*human CACCGACCAGGTATCAAGCTCATAG-3’), *KAT7* human AAACCTATGAGCTTGATACCTGGTC-3’), *BRD9* human CACCGAGATACCGTGTACTACAAGT-3’), *BRD9* human anti-sense #1 (5’-AAACACTTGTAGTACACGGTATCTC-3’), *BRD9* human sense #2 (5’-CACCGAGAGAGGGAGCACTGTGACA-3’), *BRD9* human anti-sense #2 (5’-AAACTGTCACAGTGCTCCCTCTCTC-3’), *BICRA* human sense #1 (5’-CACCG TCACCATCCAGGGCGAGCCG-3’), *BICRA* human anti-sense #1 (5’-AAACCGGCTCGCCCTGGATGGTGAC-3’), *BICRA* human sense #2 (5’-CACCGCCCGCGGGCGCAAACGGCTT-3’), *BICRA* human anti-sense #2 (5’-AAACAAGCCGTTTGCGCCCGCGGGC-3’), *CONTROL (*Non-targeting sgRNA) sense #1 (5’-CACCGAAACGGTACGACAGCGTGTG-3’), *CONTROL (*Non-targeting sgRNA) anti-sense #1 (5’-AAACCACACGCTGTCGTACCGTTTC-3’), *CONTROL (*Non-targeting sgRNA) sense #2 (5’-CACCGGTGCGCATGGGCTGATGTTA-3’), *CONTROL (*Non-targeting sgRNA) anti-sense #2 (5’-AAACTAACATCAGCCCATGCGCACC-3’). The following sgRNA oligos were used for LentiCRISPR V2-Neo vector for CRISPR knockout experiments: *BICRAL* human sense #1 (5’-CACCGACACCTTTGAGGGATGACTT-3’), *BICRAL* human anti-sense #1 (5’-AAACAAGTCATCCCTCAAAGGTGTC-3’), *BICRAL* human sense #2 (5’-CACCGCTTGGAGAAGGGCCCAGTGA-3’), *BICRAL* human anti-sense #2 (5’-AAACTCACTGGGCCCTTCTCCAAGC-3’) and sgControl #1 described above.

### Lentivirus Production

Lentiviruses were made by Lipofectamine 2000-based co-transfection of 293FT cells with the respective lentiviral expression vectors and the packaging plasmids psPAX2 (Addgene #12260) and pMD2.G (Addgene #12259) in a ratio of 4:3:1. Virus-containing supernatant was collected at 48 and 72h after transfection, pooled together (15 mL total per 10-cm tissue culture dish), passed through a 0.45-µm filter, aliquoted, and frozen at −80°C until use.

### Lentiviral Infection

For single knock-out, cells were counted using a Vi-Cell XR Cell Counter (Beckman Coulter) and 2 X10^6^ cells were resuspended in 1 mL lentivirus with 8 μg/mL polybrene in individual wells of a 12 well plate. For double knock out experiments, 4 X10^6^ cells were infected using 1 mL of mixed 1:1 sgRNA #1 and #2 targeting BICRA (puromycin resistant) and/or 1 mL of mixed 1:1 sgRNA #1 and #2 targeting BICRAL (G418 resistant) and/or 1 mL of the sgControl #1 guide (2 separate constructs that were puromycin or G418 resistant) with 8 μg/mL polybrene in individual wells of a 6 well plate. The plates were then centrifuged at 2000 rpm (931 x g) for 2h at 30°C (Allegra® X-15R Centrifuge (Beckman Coulter), rotor SX4750A). 16 hours later the virus was removed and cells were grown for 72 hours before being placed under drug selection. Cells were selected in puromycin (0.25 μg/mL) or G418 (500 μg/mL).

### Correlation Analyses of Validated Enriched Screen Hits with POU2F3 Dependency or POU2F3 Expression

Correlation of SMARCD1, BRD9, EP300, MED19, IPPK, and KAT7 dependencies with POU2F3 dependency or POU2F3 expression was analyzed using gene effect from publicly available data from Depmap (DepMap Public 23Q2+Score, Chronos)^40^. For Fig. 2, all POU2F3- positive SCLC cells (NCI-H1048, NCI-H211, NCI-H526, and CORL-311) were compared to all other SCLCs (n=23 SCLC cell lines in total with 4 POU2F3-positive SCLCs and 19 other SCLCs), and then compared to all other cancer cell lines in the dependency map. For Supplemental Fig. 2, only SCLC cell lines were included (n=23) and were binned by POU2F3 expression as above.

### SMARCD1, BRD9 and BAF complexes gene effect calculations

DepMap Public 23Q2 CRISPR based gene effect estimates for all models in the Achilles pipeline, integrated using Harmonia, were downloaded from the depmap portal (https://depmap.org/portal/). POU2F3-positive SCLCs (n=4) were compared to all other SCLCs (n=19), and then compared to all other cancer cell lines. Gene effect for each BAF complex, in Fig. 2E, was calculated based on BAF subunits reported in Michel et al., NCB, 2018^39^ (see Supplementary Table 4 for gene list). If paralogs were present in the complex, their gene effects were first averaged then the paralog average was included before averaging with all other unique subunits in the complex.

### Pharmacological Inhibitors

The following chemicals (stored at −20°C or −80°C) were added to cell culture where indicated: BVdU was purchased from Chem-Impex International Inc., catalog no. 27735. FHD-609, FHD-286, AU-15330, and dTAG-V1 were all resynthesized by Dr. Jun Qi’s laboratory according to the literature. XHC-640, WA-68-VQ71, dBRD9A, BRM014 were all synthesized by Novartis and obtained under an MTA. For immunoblot analysis, the cells were treated with the drugs at the concentrations and times indicated. For dose response assays, the cells were treated with drugs at the indicated concentrations for 6 days. For dTAG-V1 experiments, the cells were treated at 100 nM overnight. For drug treatment studies, NCI-H1048, NCI-H211, COR-L311, and NCI-H526 were seeded at 100,000 cells per mL and were treated with either DMSO, 100 nM FHD-286, or 100nM FHD-609 for 72 hours. Cells were then harvested for downstream analyses.

### Cell Proliferation Assays

For proliferation assays with NCI-H1048 SCLC CRISPR-inactivated cells in Fig. 2, cells were counted on day 0 using a Vi-Cell XR Cell Counter and plated in a tissue culture-treated 6-well plate at 15,000 cells/mL for all experiments (or 30,000 cells/mL for Fig. 2H) in 2 mL of complete media. Cells were trypsinized 3 or 6 days later to make single cell suspensions and were counted using a Vi-Cell XR Cell Counter (Beckman Coulter). Cell counts were then normalized to the number of cells plated at day 0.

For long term proliferation experiments with human POU2F3-positive SCLC cell lines in Supplemental Fig. 3, cells were counted on day 0 using a Vi-Cell XR Cell Counter (Beckman Coulter) and plated in a tissue culture-treated 6-well plates at 20,000 cells/mL in 2 mL of complete media containing XHC-640 (100 nM), WA-68-VQ71 (100 nM), dBRD9A (100 nM), or DMSO. Cells counts were performed every 3 days using the Vi-Cell until day 12. Fresh drug in fresh complete media was replaced every 3 days.

### Dose Response Assays

For dose-response assays, cells were counted as described above and plated in a tissue culture-treated 6-well plates at 20,000 cells/mL in 2 mL of complete media. For dose-response experiments performed in ASCL1- and NEUROD1-expressing cell lines, cells were counted as described above and plated in a tissue culture-treated 6-well plates at 50,000 cells/mL in 2 mL of complete media. For all dose response assays, cells were treated with the indicated drugs at 0 nM, 0.1 nM, 1 nM, 10 nM, 100 nM, 1000 nM and after 6 days were trypsinized to make single cell suspensions and counted. Cell counts were then normalized to the DMSO condition. EC50’s were calculated using non-linear regression log(inhibitor) vs. response -- Variable slope (four parameters).

### Immunoblotting

Cell pellets were lysed in a modified EBC lysis buffer (50mM Tris-Cl pH 8.0, 250 mM NaCl, 0.5% NP-40, 5 mM EDTA) supplemented with a protease inhibitor cocktail (Complete, Roche Applied Science, #11836153001) and phosphatase inhibitors (PhosSTOP Sigma #04906837001). Soluble cell extracts were quantified using the Bradford Protein Assay. 20 µg of protein per sample was boiled after adding 3X sample buffer (6.7% SDS, 33% Glycerol, 300 mM DTT, and Bromophenol Blue) to a final concentration of 1X, resolved by SDS-PAGE using either 10% or 8% SDS-PAGE, semi-dry transferred onto nitrocellulose membranes, blocked in 5% milk in Tris-Buffered Saline with 0.1%Tween 20 (TBS-T) for 1h, and probed with the indicated primary antibodies overnight at 4°C. Membranes were then washed three times in TBS-T, probed with the indicated horseradish peroxidase conjugated (HRP) secondary antibodies for 1h at room temperature, and washed three times in TBS-T. Bound antibodies were detected with enhanced chemiluminescence (ECL) western blotting detection reagents (Immobilon, Thermo Fisher Scientific, #WBKLS0500) or Supersignal West Pico (Thermo Fisher Scientific, #PI34078). The primary antibodies and dilutions used were: Rabbit Anti-POU2F3 (Cell Signaling #92579S, 1:1000), Rabbit Anti-ASCL1 (Abcam #Ab211327, 1:1000), Rabbit Anti-NEUROD1 (EPR4008) (Abcam #Ab109224, 1:1000), rabbit anti-DCK (Abcam #Ab151966, 1:2000), Rabbit anti-BRD9 (E9R2I) (Cell Signaling #58906, 1:1000), Rabbit Anti-BRD7 (D9K2T) (Cell Signaling #15125, 1:1000), Rabbit Anti-SMARCD1 (E7W9W) (Cell Signaling #35070, 1:1000), Rabbit Anti-BICRA (E6I3A) (Cell Signaling #45441, 1:1000), Rabbit Anti-BICRAL (GLTSCR1L) (Thermo Fisher Scientific #PA5-56126, 1:1000), Rabbit Anti-OCA-T1 (Cell Signaling #20217, 1:1000), Rabbit Anti-SMARCA2 (BRM) (D9E8B) (Cell Signaling #1966, 1:1000), Rabbit Anti-SMARCA4 (BRG1) (D1Q7F) (Cell Signaling #49360, 1:1000), Rabbit Anti-HA (C29F4) (Cell Signaling #3724T, 1:1000) or Mouse Anti-HA.11 (16B12) (Biolegend #901533), Rabbit Anti-GSPT1 (Abcam #Ab49878, 1:1000), Rabbit Anti-ACTL6B (E5X8C) (Cell Signaling #46787S, 1:1000), mouse α-INSM1 (Santa Cruz, #SC377428, 1:1000), Rabbit Anti-Synaptophysin (YE269) (Abcam #ab32127), and Mouse Anti-β-actin (Sigma, clone AC-15, #A3854, 1:25,000). The secondary antibodies and dilutions were: Goat Anti-Mouse (Jackson ImmunoResearch #115-035-003) and Goat anti-Rabbit (Jackson ImmunoResearch #111-035-003) and used at 1:5000.

### Human POU2F3 RNA-seq Analyses from publicly available RNA-seq Data

FPKM values from all tumors from George et al.^4^ were analyzed across all tumors (n=81) to first select tumors that highly expressed POU2F3 (high=FPKM values >2) which contained 12 tumors. 2 POU2F3-positive tumors (S01297 and S02375) had dominant expression of ASCL1 or NEUROD1 relative to POU2F3 and hence were excluded as potential confounders as ASCL1 and NEUROD1 positive tumors are associated with higher neuroendocrine gene expression.

### RNA-sequencing

For FHD-286 and FHD-609 experiments, 10 millions cells were harvested in biological triplicate and resuspended in 1mL of TRIzol Reagent (ThermoFisher). RNA was extracted following factory protocols. 5ug of RNA was then treated with DNAse to remove possible genomic DNA contamination using the DNA-free DNA Removal Kit (ThermoFisher) following factory protocols. 1ug of DNase-treated RNA was then used for NEB Next Poly(A) mRNA isolation and subsequent next generation sequencing library construction (NEB) following factory procedures. ERCC spike-in (Invitrogen) was added at a ratio of 2uL of spike-in per 1ug of RNA.

For WA-68-VQ71 and BRM014 experiments, NCI-H1048, NCI-H211, NCI-H526 and COR-L311 cells were plated at 200,000 cells/mL and treated with the small molecules indicated for 3 days in two independent biological replicates. RNA was extracted using RNeasy mini kit (Qiagen #74106) including a DNase digestion step according to the manufacturer’s instructions and RNA sequencing was performed as described below. Total RNA samples in each experiment were submitted to Novogene Inc. The libraries for RNA-seq are prepared using NEBNext Ultra II non-stranded kit. Paired end 150bp sequencing was performed on Novaseq6000 sequencer using S4 flow cell. Sequencing reads were mapped to the hg38 genome by STAR. Statistics for differentially expressed genes were calculated by DESeq2.

### ATAC-sequencing

100,000 cells were used for the OMNI-ATAC protocol^51^. Cells harvested and subsequently washed once with room temperature PBS and ice-cold PBS. Cell pellets were lysed in 50 μL cold resuspension buffer (10 mM Tris-HCl pH 7.5, 10 mM NaCl, and 3 mM MgCl_2_) supplemented at a final concentration of 0.1% NP40, 0.1% Tween-20, and 0.01% Digitonin for 3-5 minutes. Lysis step was quenched with 1mL of resuspension buffer supplemented at a final concentration of 0.1% Tween-20 and nuclei were pelleted at 500 g for 10 min at 4°C. Nuclei were then resuspended in 50 μL transposition reaction mix containing 25 μL 2X Tagment DNA buffer (Illumina), 2.5 μL Tn5 transposase (Illumina), 16.5 μL 1X PBS, 0.5 μL 1% Digitonin (final 0.01% v/v), 0.5 μL 10% Tween-20 (final 0.1% v/v), and 5 μL nuclease-free water. The transposition reaction was incubated at 37°C for 30 min with constant shaking (1000 rpm) on a thermomixer. Tagmented DNA was purified using the MinElute Reaction Cleanup Kit (Qiagen).

### ChIP-sequencing studies

In brief, cells were crosslinked at 0.8% PFA for 10 minutes at room temperature. After crosslinking reactions were quenched with 150mM glycine for 10 minutes at room temperatures. Cells were washed twice with ice cold PBS and then either flash frozen in liquid nitrogen or resuspended in cell lysis buffer (50mM HEPES pH7.5, 90mM KCl, 1mM EDTA, 10% Glycerol, 0.5% NP-40, 0.25% Triton X-100) for 15 minutes. Cell mixtures were dounced with a tight fitting glass dounce with 20-30 strokes. Nuclei were washed twice with ice cold MNase Buffer (50mM HEPES pH7.5, 1mM CaCl2, 20mM NaCl). The nuclei were then resuspended at ∼10-20 million nuclei per mL of sonication buffer (50mM, 1mM CaCl2, 140mM NaCl, 5mM EDTA, 2.5mM EGTA, 0.1% sodium deoxycholate, 0.4% sodium lauroyl sarcosinate) and added into 1mL milliFiber Covaris tubes. Sonication was performed at 140 PIP, 6.0% DF, and 200 CPB at a rate of 60 seconds ON/ 30 seconds off for 10-16 cycles depending on the cell line. After sonication, sonicated chromatin was supplemented to final concentration of 1.1% Triton-X and 5% glycerol and subsequently set up for immunoprecipitation with the following antibodies: Rabbit Anti-POU2F3 (Cell Signaling E5N2D, #36135, 1:50), mouse anti-Rpb1 (Cell Signaling 4H8, #2629, 1:50), Rabbit Anti-H3K27Ac (Cell Signaling D5E4, #8173, 1:100).

IPs were performed overnight at 4°C and then incubated with animal-specific dynabeads (ThermoFisher) for 2.5 hours at 4°C. Beads were washed 3X with RIPA150 (10mM Tris HCL pH7.5, 1mM EDTA, 0.1% sodium dodecyl sulfate, 0.1% sodium deoxycholate, 1% Triton X-100, 150mM NaCl), then 3X with RIPA500 (10mM Tris HCL pH7.5, 1mM EDTA, 0.1% sodium dodecyl sulfate, 0.1% sodium deoxycholate, 1% Triton X-100, 500mM NaCl), and 3X with LiCl Wash Buffer (250mM LiCl, 0.5% NP-40, 0.5% sodium deoxycholate). Beads were then eluted with ChIP Elution Buffer (50mM Tris HCL pH 7.5, 10mM EDTA, 1% sodium dodecyl sulfate) and reverse crosslinked with proteinase K overnight at 65C and subsequently treated with RNAse A (NEB) for 1 hour at 37°C. ChIP DNA was purified using Qiagen MinElute Reaction Cleanup kit (Qiagen) following factory protocols.

### Reverse-Transcriptase Quantitative PCR (RT-qPCR)

RNA was extracted using Quick-RNA™ Miniprep kit (Zymo Research, CA, USA) according to the manufacturer’s instructions. RNA concentration was determined using the Nanodrop 8000 (Thermofisher Scientific). A cDNA library was synthesized using iScript Reverse Transcription Supermix for RT-qPCR (Biorad #1708841) according to the manufacturer’s instructions. qPCR for *COLCA2* (OCA-T2) was performed using the LightCycler 480 (Roche) with the LightCycler 480 Probes Master Kit (Roche) and Taqman probes (ThermoFisher Scientific) according to the manufacturer’s instructions. The Λ1Λ1C_T_ Method was used to analyze data. The C_T_ values for each probe were then normalized to the C_T_ value of *ActB*. The following TaqMan probes were used: *ACTB* human (Hs01060665_g1) and *COLCA2 (*OCA-T2) human (Hs00416978_m1).

### Data processing for RNA-seq, ATAC-seq, and ChIP-Seq

RNA-seq and ATAC-seq samples were demultiplexed using bcl2fastq v2.20.0.422. The sequenced RNA-seq reads were aligned to hg19 assembly with STAR v2.5.2b^52^. The sequenced ATAC-seq reads were first trimmed with Trimmomatic v0.36, aligned with Bowtie2 v2.2.9, and filtered with Picard v2.8.0 (MarkDuplicates REMOVE_DUPLICATES=true) and SAMtools v 0.1.19 (-F 256 -f 2 -q 30)^53–55^. Reads mapping to regions defined in the ENCODE project’s wgEncodeDacMapabilityConcensusExcludeable bed file were removed using bedtools v2.30.0^56^. The sequenced ChIP-seq reads were processed identically to ATAC-seq reads. All genomics data (ATAC-seq, RNA-seq, and ChIP-seq) have been deposited to the Gene Expression Omnibus (GEO) under accession number: GSE249362 except RNA-seq data of cells treated with BRM014 and WA-68-VQ71 are in GEO under accession number GSE249258. Data will be released upon publication.

### ATAC-seq analysis

ATAC reads were merged across technical replicates using samtools merge^53^. Reads were filtered to only include reads with an insert size of 38-100bp, pre-shift, using htslib^57^. Peaks were called using MACS3 callpeak (-q 0.01 --nomodel --extsize 200)^58^. Venn diagrams of peak overlaps were generated using bedtools intersect, and visualized using the eulerr R package^56,59^. PCA plots for ATAC were plotted using the top 500 peaks by variance. Counts were determined using bedtools intersect, then variance stabilizing transformation was applied. FASTA sequences across given sites were generated using site centers with flanking windows of 200bp (total window size of 400bp). Enriched motifs across these sets of sites were determined using HOMER findMotifsGenome.pl against genome-background (-size 400)^60^. HOMER motif known results were visualized as barplots using ggplot2.Heatmaps and metaplots were generated over indicated peaks using deepTools computeMatrix^61^. Bigwig inputs for heatmaps were generated with deepTools bamCoverage (--binSize “40” --normalizeUsing “CPM” --exactScaling)^61^.Different sets of sites were assigned to their nearest protein-coding gene using bedtools closest, These distances were visualized as stacked bar charts using ggplot in R, highlighting the proportion of promoter, promoter proximal, and distal enhancer regions.

### ChIP-seq analysis

Bigwigs were again generated using deepTools bamCoverage (--binSize “40” -- normalizeUsing “CPM” --exactScaling) and visualized using IGV^62^. Peaks were called using MACS3 callpeak (-q 0.01 --nomodel --extsize 200)^58^.

### RNA-seq Data Analysis

Identification of upregulated or downregulated genes across the conditions were determined using DESeq2^51^ (log2FC = 1, B-H p-value = 0.05). Normalized counts were generated using DESeq2’s estimateSizeFactors function. Volcano plots for changes in expression were visualized as scatter plots using ggplot2^63^. Venn diagrams of differential genes were generated using the eulerr R package^56,59^.

PCA plots were generated using variance stabilizing transformed counts, across the top 500 genes by variance, and plotted using ggplot2. Heatmaps were generated using the ComplexHeatmap R package^64^, and visualize Z-Score normalized read counts for each gene across all samples for each cell line, unless otherwise indicated. Hierarchical clustering was performed using Spearman or Kendall distance functions.

GSEA analysis was performed using the Hallmark gene sets, through the msigdbr R package, using clusterProfiler GSEA function^65^. Differential expression ranking was determined by the “stat” output from DESeq2. For GSEA analysis in Fig. 6A-B, S6A-B, GSEA software was obtained from the Gene Set Enrichment Analysis website [http://www.broad.mit.edu/gsea/downloads.jsp]. Top 200 downregulated gene lists were identified as significantly downregulated genes (adjusted p value <0.05) with top 200 log transformed fold decrease after FHD-286 or BRM014 treatment (100 nM for 72 hours) on NCI-H211, NCI-H1048, NCI-H526, and COR-L311, FHD-609 or WA-68-VQ71 treatment (100 nM for 72 hours) on NCI-H211 and NCI-H1048. Subsequently, pre-ranked GSEA was performed on log transformed fold change profiles from RNA-sequencing data reported in Wu et al., Nature, 2022^13^, where POU2F3, OCA-T1 or OCA-T2 was knocked out in NCI-H211, NCI-H1048, NCI-H526, and COR-L311 cell lines.

For Figure 4H, ATAC fold-changes were determined by finding ATAC peaks in both conditions within +/− 10kb of the differentially expressed gene’s TSS, and calculating the fold change in reads-per-million across all such peaks.

For Figure 4M, primary targets were identified for differentially expressed genes whose TSS were within +/− 10kb of a concordant change in accessibility, using bedtools closest and the GENCODE consortium version 44 annotation for GRCh37.

### Exogenous POU2F3 and OCA-T2 Rescue Experiments

For HA-dTAG-POU2F3 or OCA-T2-dTAG-HA system, the FKBP23F36V-2xHA was PCR amplified from the pCRIS-PITCHv2-Puro-dTAG vector (addgene #91703) and introduced into sgRNA-resistant POU2F3_LentiV_NEO or the OCA-T2_LentiV_neo vector for functional validation with competition-based cell proliferation assay. NCI-H1048 cells that stably expressed Cas9 were infected either with HA_dTAG_POU2F3_LentiV_neo or OCA-T2_dTAG_HA_LentiV_neo or empty_vector_lentiV_neo construct followed by neomycin selection to establish stable cell lines. The cells were then lentivirally delivered with indicated sgRNAs co-expressed with a GFP reporter. The percentage of GFP+ cells corresponding to the sgRNA representation within the population. GFP measurements in human cell lines were taken on day 4 post-infection and every four days with Guava Easycyte HT instrument (Millipore). The fold change in GFP+ population (normalized to day 4) were used for analysis. After validating the functionality of tagged POU2F3/OCA-T2, the HA_dTAG_POU2F3 or OCA-T2_dTAG_HA, which is linked with blasticidin resistant gene through P2A linker (HA_dTAG_POU2F3_P2A_blasticidin, or OCA-T2_dTAG_HA_P2A_Blasticidin) and resistant to its own sgRNA, were cloned into the LRG2.1T vector that either contains sgRNA against endogenous POU2F3 or OCA-T2 into NCI-H1048 that stably express Cas9. The following sgRNA oligos were used for CRISPR knockout : sgControl #1 = AGTCGCTTCTCGATTATGGG, sgControl #2 = CAGAGTCTCCTATGCCACAC, sgCDK1=ACACAATCCCCTGTAGGATT, sgPOU2F3 #4 = GACCAACATCCGCCTGACTC, sgOCA-T2 #1=CGGACACCTTGATACACCTT and sgOCA-T2 #2= CCGAGTGAAGATCACAGTGA.

For immunoblot analyses in NCI-H1048 cells expressing HA-dTAG-POU2F3, or HA-dTAG-OCA-T2, or parental NCI-H1048 cells, the cells were plated at 200,000 cells/mL and treated with the small molecules indicated for 3 days. For the dose response assays, the cells were plated at 20,000 cells/mL and treated with the small molecules indicated for 6 days and counted using the Vi-Cell. Cell counts were then normalized to the number of cells plated at day 0. EC50’s were calculated using non-linear regression log(inhibitor) vs. response-variable slope (four parameters).

### SCLC Cell Line Xenografts and Treatment Studies

For NCI-1048 and NCI-H211 xenografts, parental cells were grown to 10^9^ cells, washed 3 times in 50 mLs of sterile PBS, and resuspended at 6 x 10^7^ cells/mL in PBS with 50% Matrigel (Westnet Inc. #356231) for NCI-H1048 xenografts or resuspended at 10 x 10^7^ cells/mL in PBS with 50% Matrigel (Fisher Scientific #CB40234) for NCI-H211 xenografts. The mice were anesthesized with isoflurane and 6 x 10^6^ cells (for NCI-H1048) or 10 x 10^6^ cells (for NCI-H211) were injected subcutaneously into bilateral flanks of 6 week old Ncr nude female mice (Taconic #NCRNU). The mice were monitored daily and when flank tumors were ∼100 mm^3^ in size (∼2 weeks after injection), mice were randomized to treatment with either FHD-609, FHD-286, or vehicle (HP-β-CD, Sigma Aldrich #778966). Vehicle was first made by resuspending HP-β-CD at 20% in UltraPure Distilled Water (Thermo Fisher Scientific #10977023). FHD-609 and FHD-286 powder was weighed using a high precision scale (Mettler Toledo, XS105), resuspended in the HP-β-CD vehicle and rotated overnight at 4°C. All drugs, including vehicle, were made fresh twice a week and stored in the dark at 4°C. FHD-609 was dosed daily at 0.5 mg/kg by intraperitoneal (IP) injections. Vehicle was also dosed daily by IP injections. FHD-286 was dosed daily at 1.5 mg/kg by oral gavage (PO). For the PD study, mice were dosed daily for 7 days and tumors were harvested 6 hours after the last dose and flash frozen (2/3) or fixed (1/3) for downstream analysis. For the efficacy study, mice were dosed continuously for 35 days at which time they were monitored off treatment until each mouse reached its endpoint. All mice with tumors were enrolled and no data were excluded. Tumor diameters were measured twice a week using calipers until mice were euthanized and tumor volume was calculated using: tumor volume (mm^3^) = (width)^2^ x length/2. Body weights were also measured twice a week to monitor for overt toxicity. Mice were euthanized when one of the tumors reached their endpoint of >1500mm^3^. In the NCI-H211 xenograft experiment, there was 1 mouse in the FHD-286 arm and 1 mouse in the vehicle arm that had a tumor that didn’t grow following treatment completion. All mice were followed up until 70 days at which point all other mice reached their endpoint except for the 2 mice above. At this point, the study was ended and both of these mice were also euthanized. The Kaplan Meier Estimator was performed to analyze median overall survival. Upon euthanization, ∼2/3 of each lung tumor was immediately flash frozen on dry ice for subsequent RNA and protein analysis, ∼1/3 of each lung tumor was fixed in 10% formalin for 24 hours then stored in 70% ethanol before being embedded in paraffin.

### Statistics

For the *in vivo* xenograft studies, both raw tumor volumes and log tumor fold-change normalized to day 0 are shown. The tumor volume fold-changes were log-transformed to stabilize the variability across time points. Log tumor fold-changes were modeled over the duration of treatment (until day 35) using a linear mixed-effects model accounting for the repeated measures within each tumor. The model included time (days), arm (FHD-609, FHD-286, and vehicle control), and the interaction between time and arm as covariates; these interaction terms evaluated the differences in the rate of change of the log tumor fold-change over time between arms and were considered significant at the 0.10 level. Survival was evaluated using Kaplan-Meier survival curves, and Gehan-Breslow-Wilcoxon tests were used to compare survival curves between arms as treatment only occurred during the first 35 days of the study and thereafter mice were monitored off treatment for survival. p-values of <0.05 were considered statistically significant.

For screen replicate reproducibility, r Pearson correlation coefficient’s were calculated. For the positive-selection BVdU resistance CRISPR screen analysis, Apron, Hypergeometric, and STARS analyses were performed and included in Supplementary Figure 1. For all other experiments, statistical significance was calculated using unpaired, two-tailed Students t-test. *p*-values were considered statistically significant if the *p*-value was <0.05. For all figures, * indicates *p*-value <0.05, ** indicates *p*-value <0.01, *** indicates *p*-value <0.001, and **** indicates *p*-value <0.0001. Error bars represent mean +/− SEM unless otherwise indicated.

## Supplementary Tables

**Supplementary Table 1. Raw data and gene level analysis for genome-scale positive selection screen for POU2F3 regulators.** (Tab 1) Raw Counts from the screen shown in Fig. 1 in the NCI-H1048 POU2F3-DCK* cells before treatment (Day 9), or in the BVdU or Untreated arm 7 days later (Day 16) for 2 independent biological replicates. (Tab 2) Log Normalized Counts from the screen shown in Fig. 1 in the NCI-H1048 POU2F3-DCK* cells before treatment (Day 9), or in the BVdU or Untreated arm 7 days later (Day 16) for 2 independent replicates. (Tab 3) APRON analysis of the screen showing BVDU arm at Day 16 (replicates 1 and 2) compared to Untreated arm at Day 16 (replicates 1 and 2). (Tab 4) Hypergeometric analysis of the screen showing BVDU arm at Day 16 (replicates 1 and 2) compared to Untreated arm at Day 16 (replicates 1 and 2). (Tab 5) STARS analysis of the screen showing enriched hits in BVDU arm at Day 16 (replicates 1 and 2) compared to Untreated arm (replicates 1 and 2). (Tab 6) STARS ANALYSIS of the screen showing depleted hits in BVDU arm at Day 16 (replicates 1 and 2) compared to Untreated arm at Day 16 (replicates 1 and 2). (Tab 7) Scores of hits for secondary validation in Fig. 2A using Apron, Hypergeometric and STARS analyses.

**Supplementary Table 2. Expression of SCLC subtype markers ASCL1, NEUROD1, and POU2F3 in POU2F3-positive human SCLC.**

**Supplementary Table 3. Sequences used to knock-in DCK* into the endogenous POU2F3 locus of the NCI-H1048 human SCLC cell line.** gBlocks^TM^ gene fragment sequences ordered, corresponding to the Homology arm 1 (HA1) and 2 (HA2), and sequences corresponding to the linker1, linker2, DCK*, P2A, GFP, M13 pCU and M13 Rv use to knock-in the DCK*-P2A-GFP sequence before the endogenous stop codon of *POU2F3* gene in the NCI-H1048 human SCLC cell line using CRISPR/Cas9-mediated homologous recombination strategy.

**Supplementary Table 4. mSWI/SNF complexes subunits.** List of subunits used to calculate gene effect scores for each mSWI/SNF complexes (ncBAF, cBAF and PBAF) in POU2F3-positive SCLCs relative to all other SCLC cell lines or relative to all other cancer cell lines shown in Supplementary Fig. 2.

