## Supplemental Figures and Legends for "Mammalian SWI/SNF complex activity regulates POU2F3 and constitutes a targetable dependency in small cell lung cancer"

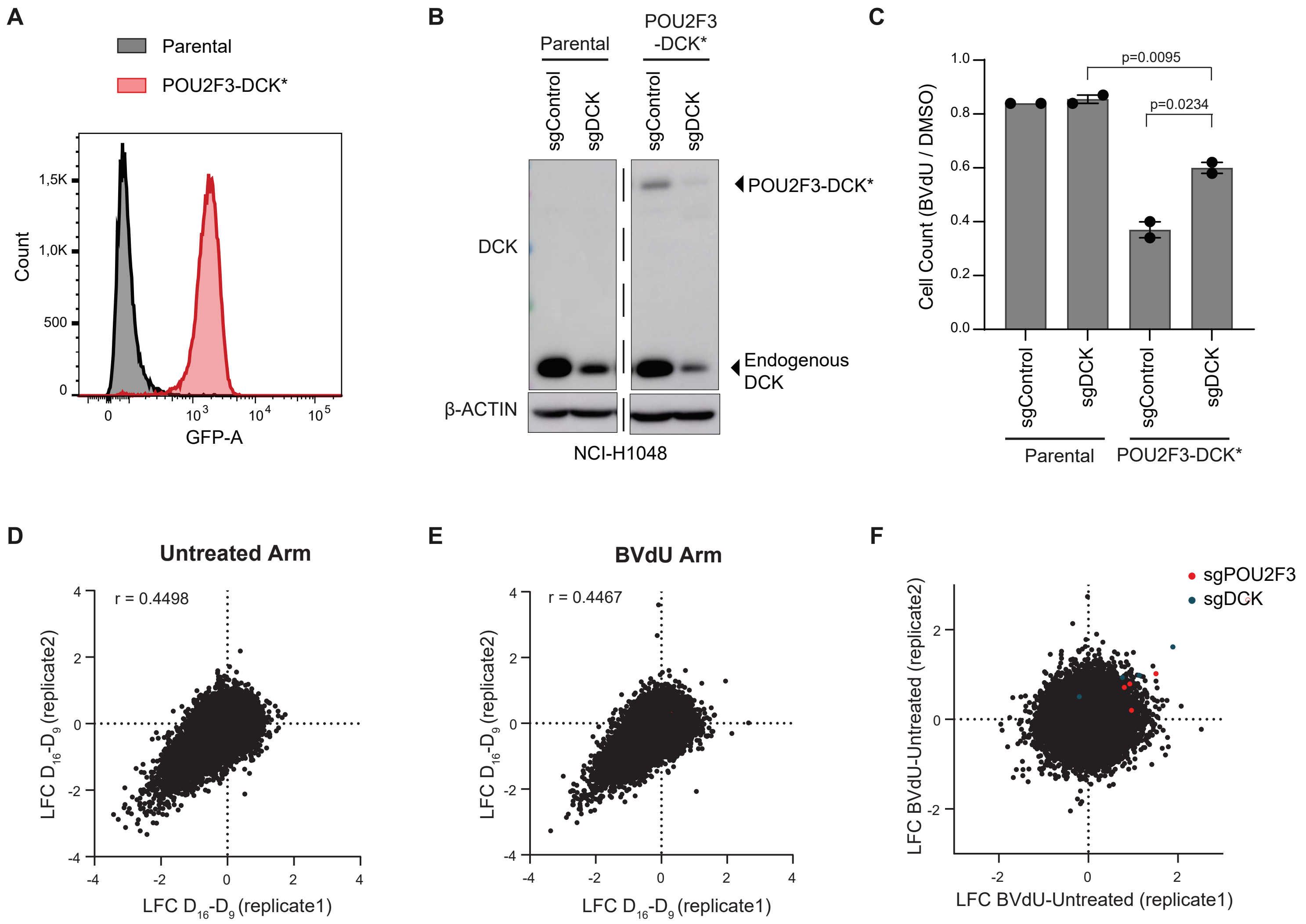


**Supplemental Fig. 1. Related to Figure 1.** (**A**) Quantification of the number of cells expressing GFP by flow cytometry in NCI-H1048 parental or NCI-H1048 POU2F3-DCK*-P2A-GFP knock-in cells. (**B**) Immunoblot analysis of NCI-H1048 parental or NCI-H1048 POU2F3-DCK*-P2A-GFP knock-in cells infected with lentiviruses encoding an sgRNA targeting DCK (sgDCK) or a non-targeting sgRNA (sgControl). (**C**) BVdU sensitivity assay of the indicated cell lines from B treated with 10μM BVdU for 7 days showing viable cell counts relative to the DMSO control. n = 2 biological replicates. p-values are indicated on figure. Error bars represent mean +/- SEM. (**D-E**) Replicate reproducibility plot of the screen show in Fig.1 showing Log Fold Change (LFC) of each individual sgRNA at day 16 compared to Day 9 in replicate 1 vs. replicate 2 of the untreated arm (D) or the BVdU-treated arm (E). For D and E, r Pearson correlation coefficient's are indicated. (**F**) Log Fold Change (LFC) of each sgRNA enriched in the BVdU arm compared to the untreated arm at day 16 in replicate 1 vs. replicate 2. sgRNAs targeting DCK are in blue and sgRNAs targeting POU2F3 are in red.

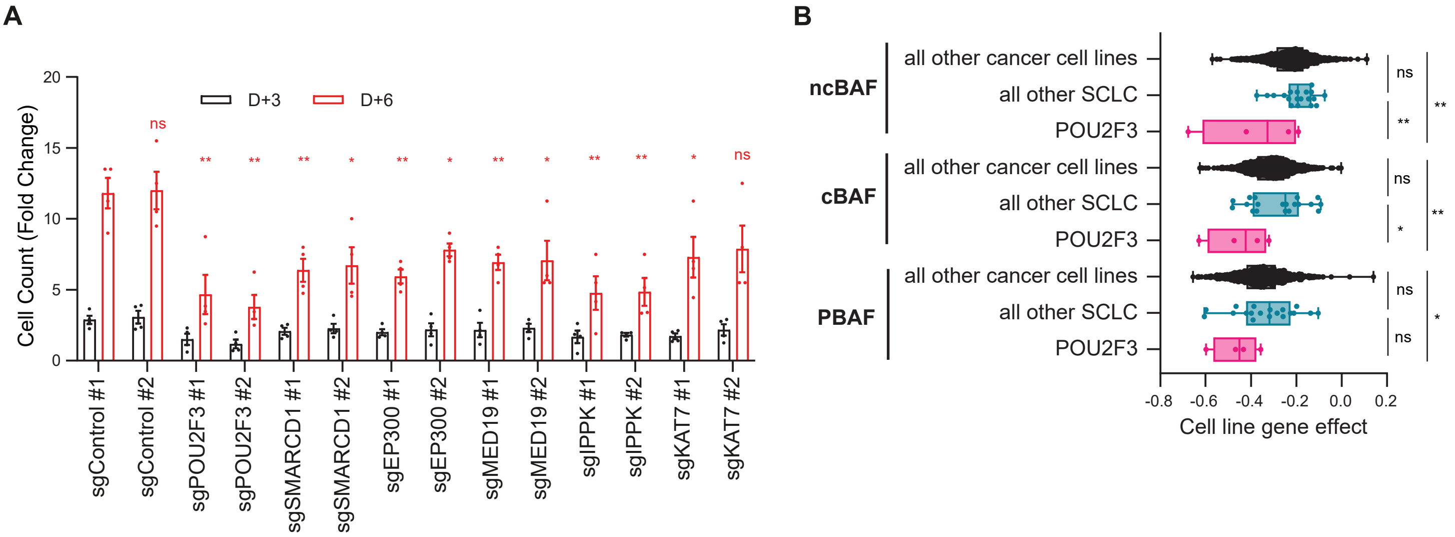


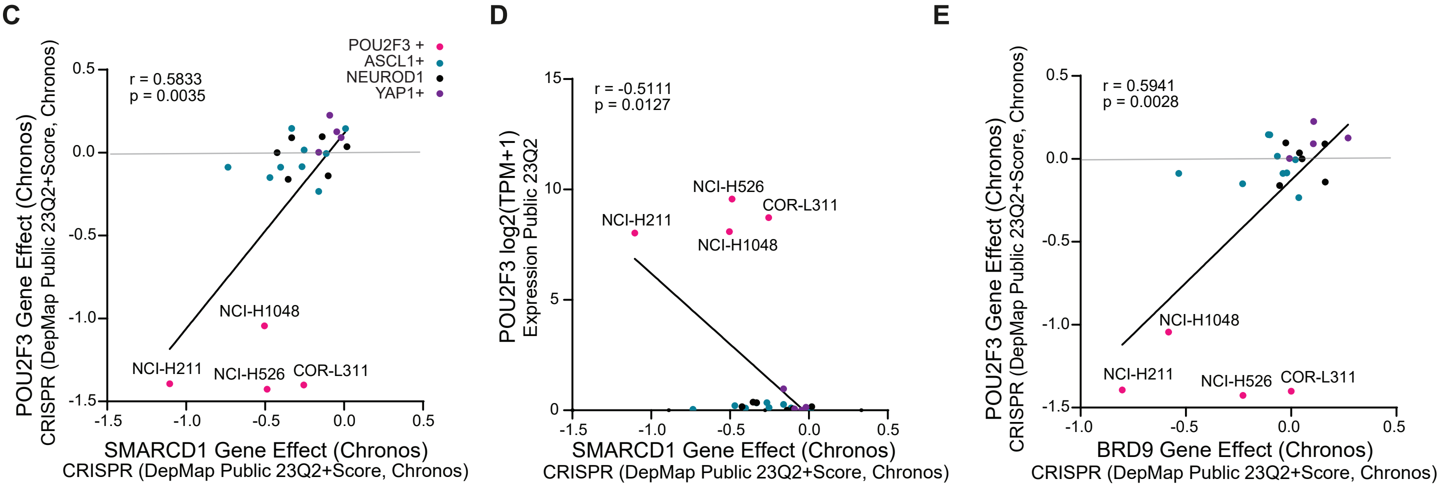


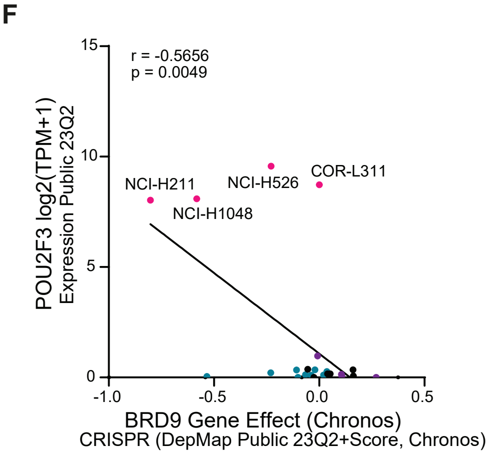


**G**

| | POU2F3 Gene Effect (Chronos) | | POU2F3 log2 (TPM+1) | |
|---|---|---|---|---|
| | r | p-value | r | p-value |
| EP300 Gene Effect (Chronos) | 0.2318 | 0.2873 | -0.2482 | 0.2534 |
| MED19 Gene Effect (Chronos) | 0.3088 | 0.1516 | -0.2294 | 0.2924 |
| IPPK Gene Effect (Chronos) | -0.1594 | 0.4676 | 0.1454 | 0.5081 |
| KAT7 Gene Effect (Chronos) | -0.1261 | 0.5665 | 0.09203 | 0.6762 |

**Supplementary Fig.2 Related to Figure 2.** (**A**) Quantitation of cell counts of NCI-H1048 cells infected with the indicated sgRNAs. Cell counts were performed 3 and 6 days after plating. Data are plotted as fold change relative to day 0. n=4 biological replicates. (**B**) Analysis of mSWI/SNF complexes (cBAF, PBAF and ncBAF) gene effect scores in POU2F3-positive SCLCs relative to all other SCLC cell lines or relative to all other cancer cell lines in the dependency map. 2-tailed unpaired t-test was used to generate p-values. n=4 POU2F3-expressing SCLC cell lines, n=19 other SCLC cell lines, n=1072 all other cancer cell lines. (**C-F**) Correlation of SMARCD1 dependency (C, D) or BRD9 dependency (E, F) vs. POU2F3 dependency (C,E) or POU2F3 expression (D,F) across SCLC cell lines. r=Pearson correlation coefficient. p-value was calculated using a two-sided Pearson's correlation test. n=4 POU2F3-expressing SCLC cell lines, n=9 ASCL1-expressing SCLC cell lines, n=6 NEUROD1-expressing SCLC cell lines, n=4 YAP1-expressing SCLC cell lines. (**G**) Correlation of EP300, MED19, IPPK or KAT7 dependency vs. POU2F3 dependency (left) or POU2F3 expression (right). r=Pearson correlation coefficient. p-value was calculated using a two-sided Pearson's correlation test.

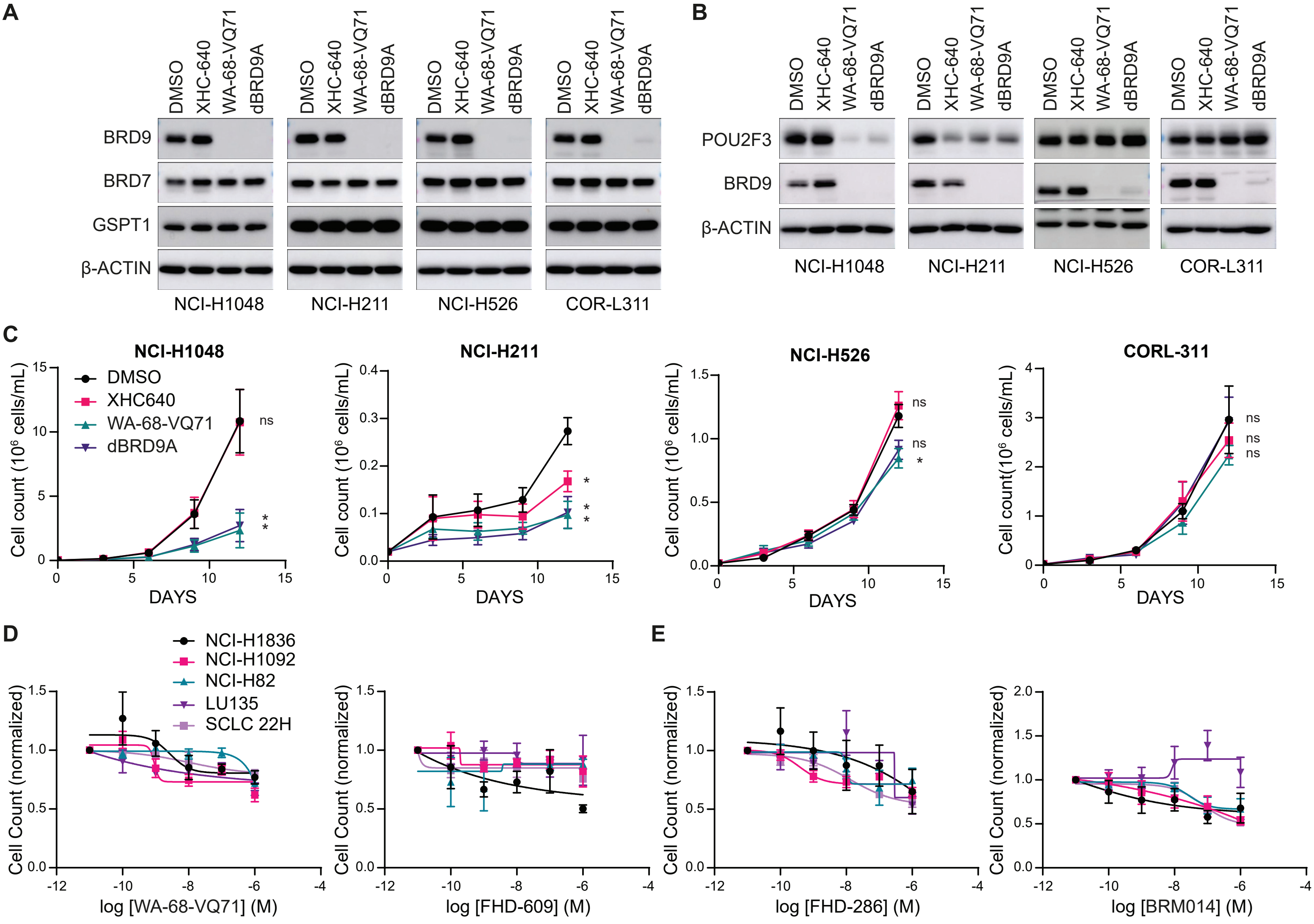


**F**

**EC50 (nM)**

| | | WA-68-VQ71 | FHD-609 | FHD-286 | BRM014 |
|---|---|---|---|---|---|
| ASCL1+ | NCI-H1836 | >1000 | =1000 | >1000 | >1000 |
| | NCI-H1092 | >1000 | >1000 | >1000 | >1000 |
| NEUROD1+ | NCI-H82 | >1000 | >1000 | >1000 | >1000 |
| | LU135 | >1000 | >1000 | >1000 | >1000 |
| Mixed ASCL1+/NEUROD1+ | SCLC 22H | >1000 | >1000 | >1000 | >1000 |

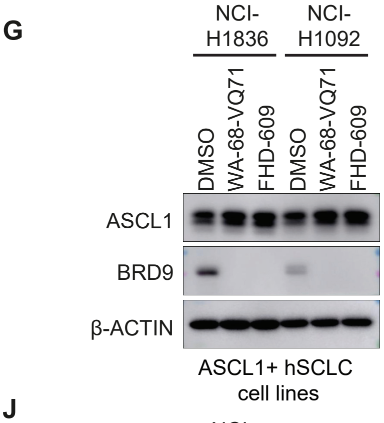


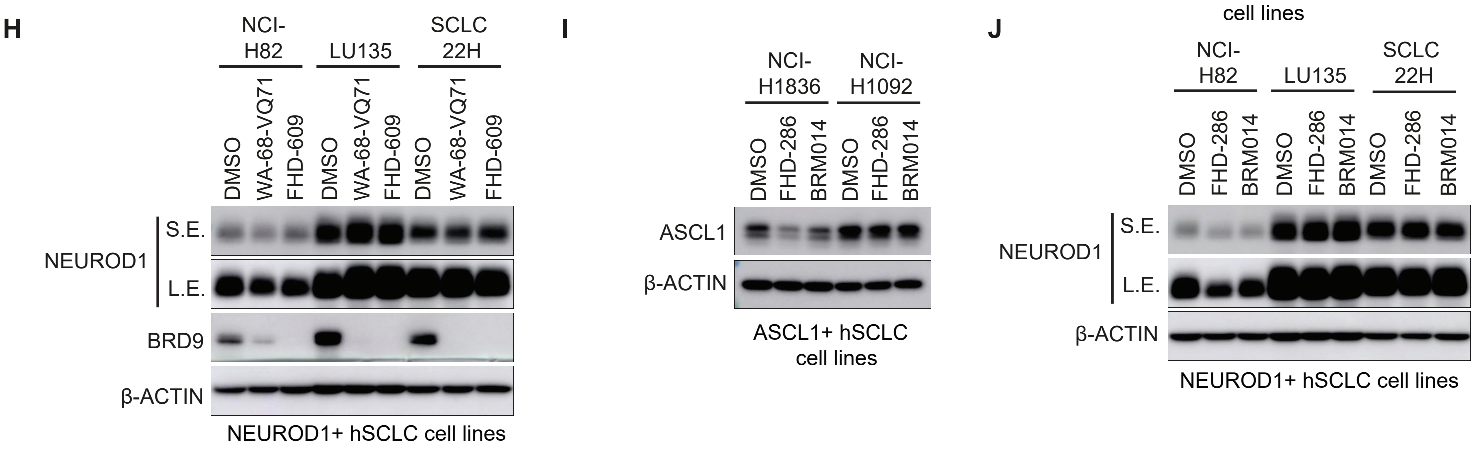


**Supplementary Fig. 3. Related to Figure 3.** (**A-B**) Immunoblot analysis of NCI-H1048, NCI-H211, NCI-526 and COR-L311 POU2F3-positive human SCLC cell lines after overnight treatment (A) or 3 days of treatment (B) with the BRD9 inhibitor (XHC-640) or 2 different BRD9 degraders (WA-68-VQ71 or dBRD9A) at 100nM or DMSO. (**C**) Proliferation assays of NCI-H1048, NCI-H211, NCI-526, and COR-L311 cells treated with a BRD9 inhibitor (XHC-640) or 2 different BRD9 degraders (WA-68-VQ71 or dBRD9A) at 100nM or DMSO for 12 days. Counts were performed every 3 days. n=3 biological replicates. *=p<0.05 using a 2-tailed unpaired t-test. p-values were calculated on day 12 counts. Error bars represent mean +/- SEM. (**D-E**) Dose response assays on the indicated ASCL1- or NEUROD1-expressing human SCLC cell lines treated for 6 days with 2 different BRD9 degraders; FHD-609 or WA-68-VQ71 (D); or 2 different SMARCA2/4 inhibitors; FHD-286, BRM014 (E). (**F**) Table showing calculated EC50's from the dose response curves from D,E. For D,E, n=3 biological replicates. (**G-J**) Immunoblot analysis on the indicated ASCL1-positive (G, I), NEUROD1-positive or mixed ASCL1+/NEUROD1-positive (H, J) human SCLC cell lines treated for 3 days with 2 different BRD9 degraders (WA-68-VQ71 and FHD-609) (G,H), or 2 different SMARCA2/4 inhibitors (FHD-286 and BRM014) (I,J).

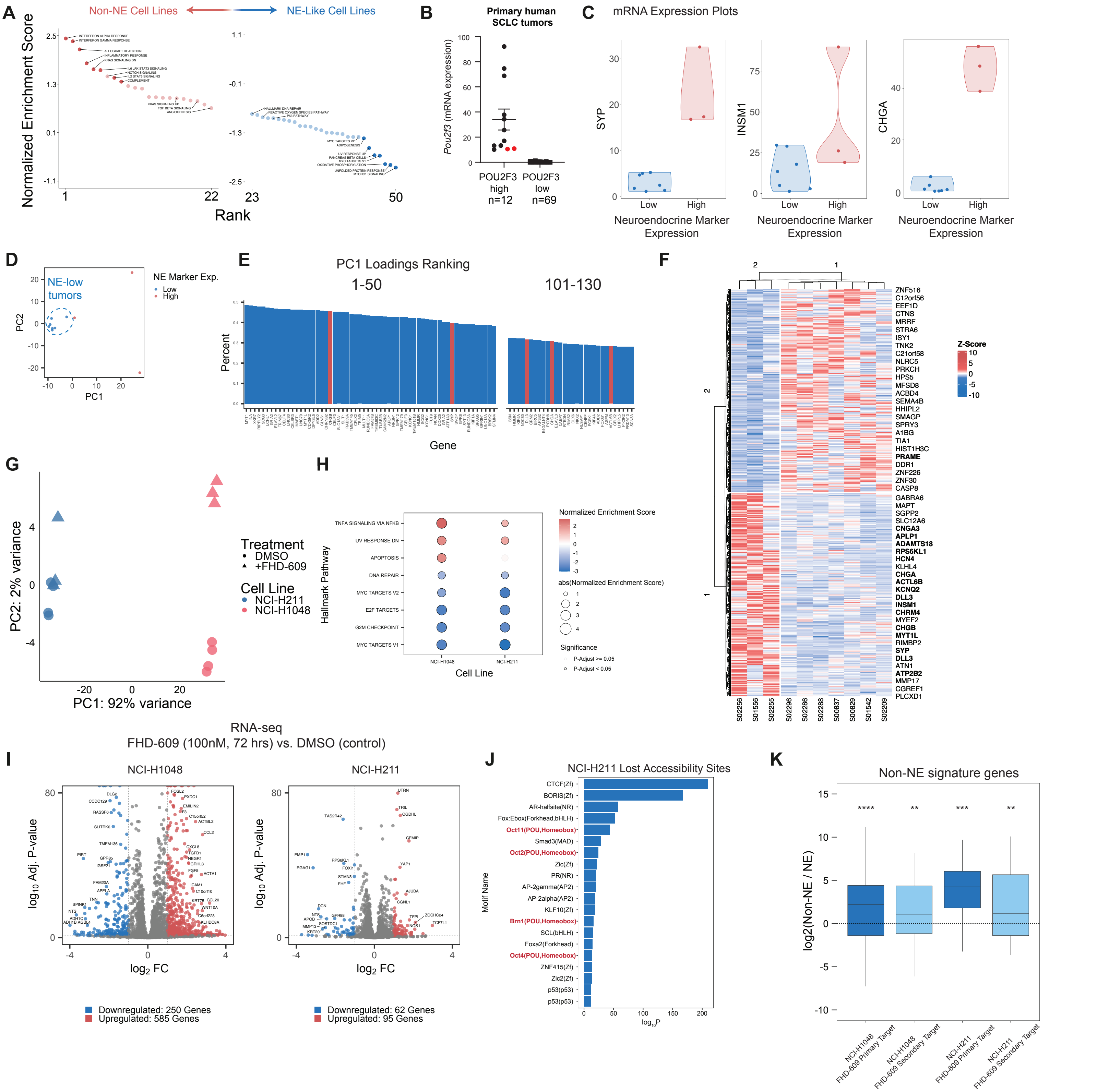


**Figure S4, Related to Figure 4. Gene expression signatures and response to pharmacologic ncBAF disruption in SCLC cell lines and primary tumors.** (**A**) Normalized enrichment scores for GSEA pathways in non-NE and NE-like cell lines. (**B**) POU2F3 expression across n=81 primary SCLC tumors where n=12 express POU2F3 and n=69 have no POU2F3 expression. POU2F3-positive tumors marked in red (S01297 and S02375) had dominant expression of ASCL1 or NEUROD1 relative to POU2F3 and so were excluded for subsequent analyses. (**C**) mRNA expression plots for SYP, INSM1, and CHGA. (**D**) PCA performed on primary tumors showing separation between NE-like and non-NE. (**E**) Loadings (genes) driving PC1 in the PCA in (c). (**F**) Gene expression in n=10 primary SCLC tumors with POU2F3 expression. Genes highlighted in bold are those from the Balanis et al., 2019 NE SCLC signature. (**G**) PCA for NCI-H1048 and NCI-H211 cell lines upon FHD-609 treatment. (**H**) Top up- and down-regulated pathways in NCI-H1048 and NCI-H211 cell lines upon FHD-609 treatment. (**I**) Volcano plots depicting up- and down regulated genes upon FHD-609 treatment; significant DEGs are highlighted in red and blue, selected genes labeled. (**J**) HOMER motif analysis performed over sites with lost accessibility upon FHD-609 treatment in NCI-H211 cells. (**K**) Non-NE signature genes are enriched in primary target genes identified by integrative ATAC- and RNA-seq analyses in non-NE SCLC cell lines treated with FHD-609.

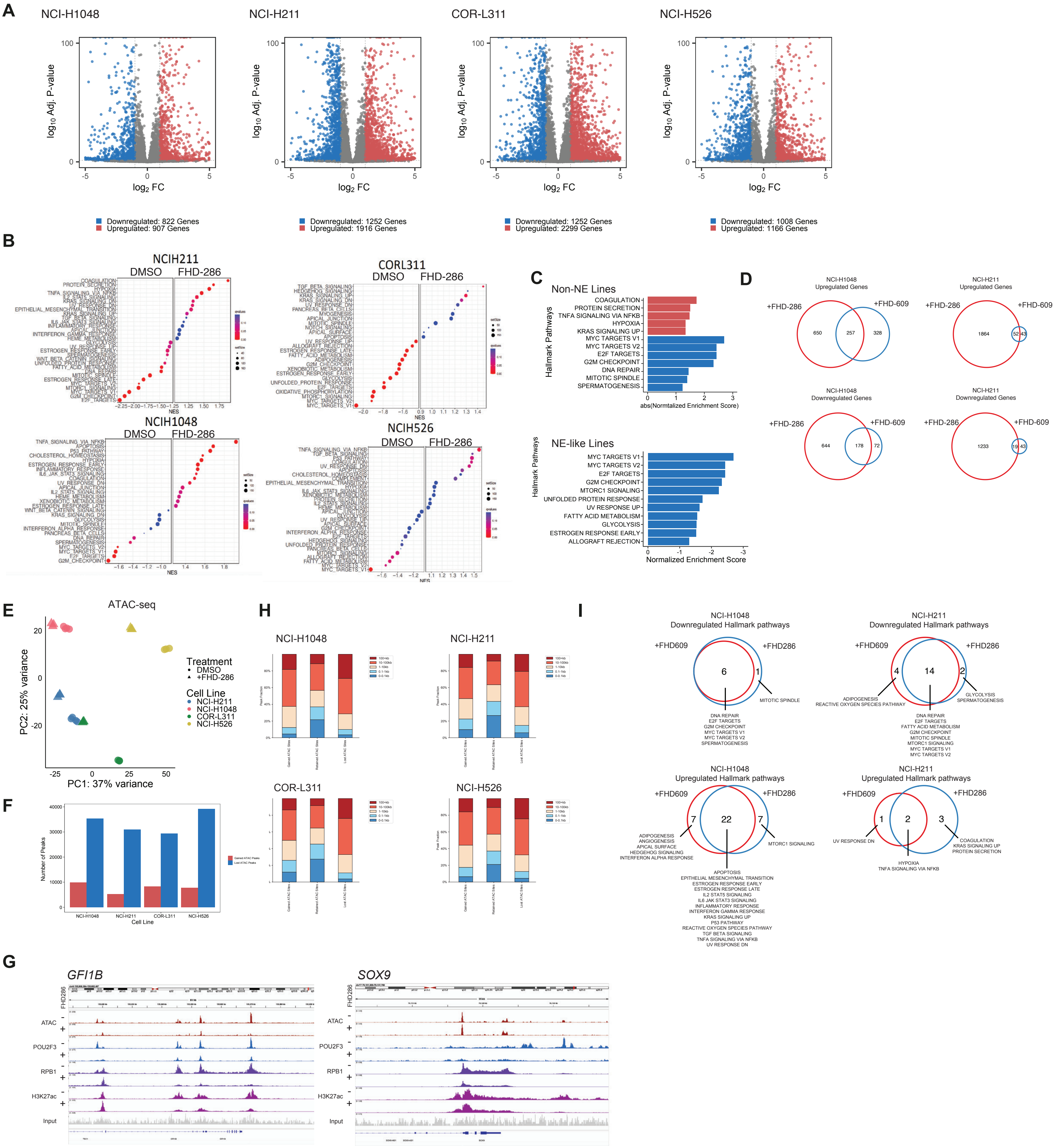


**Figure S5, related to Figure 5. mSWI/SNF disruption-specific and shared changes in gene and chromatin regulation in SCLC.** (**A**) Volcano plots showing significantly down- (blue) and up- (red) regulated genes across all cell lines upon treatment with FHD-286. (**B**) Dot plots showing gene expression pathways up- and down-regulated by treatment with FHD-286. (**C**) GSEA analyses performed on merged upregulate and downregulated genes in non-NE and NE-like cell lines. (**D**) Venn diagrams depicting shared and treatment-specific up- and down-regulated genes in each cell line treated with FHD-286 and FHD-609. (**E**) PCA performed on ATAC-seq peaks defined in all four SCLC cell lines treated with either DMSO control or FHD-286. (**F**) Bar graph depicting number of lost (blue) and gained (red) peaks in ATAC-seq experiments performed across all four cell lines. (**G**) Representative tracks at the GFI1B and SOX9 loci showing ATAC-seq signal, and POU2F3, RBP1, H3K27Ac ChIP-seq in DMSO and FHD-286 conditions. (**H**) Stacked bar graphs showing distance-to-TSS across lost, retained, and gained ATAC-seq sites upon FHD-286 treatment in each cell line. (**I**) Venn diagrams showing shared and treatment-specific changes in gene expression pathways upon FHD-286 and FHD-609 treatments.

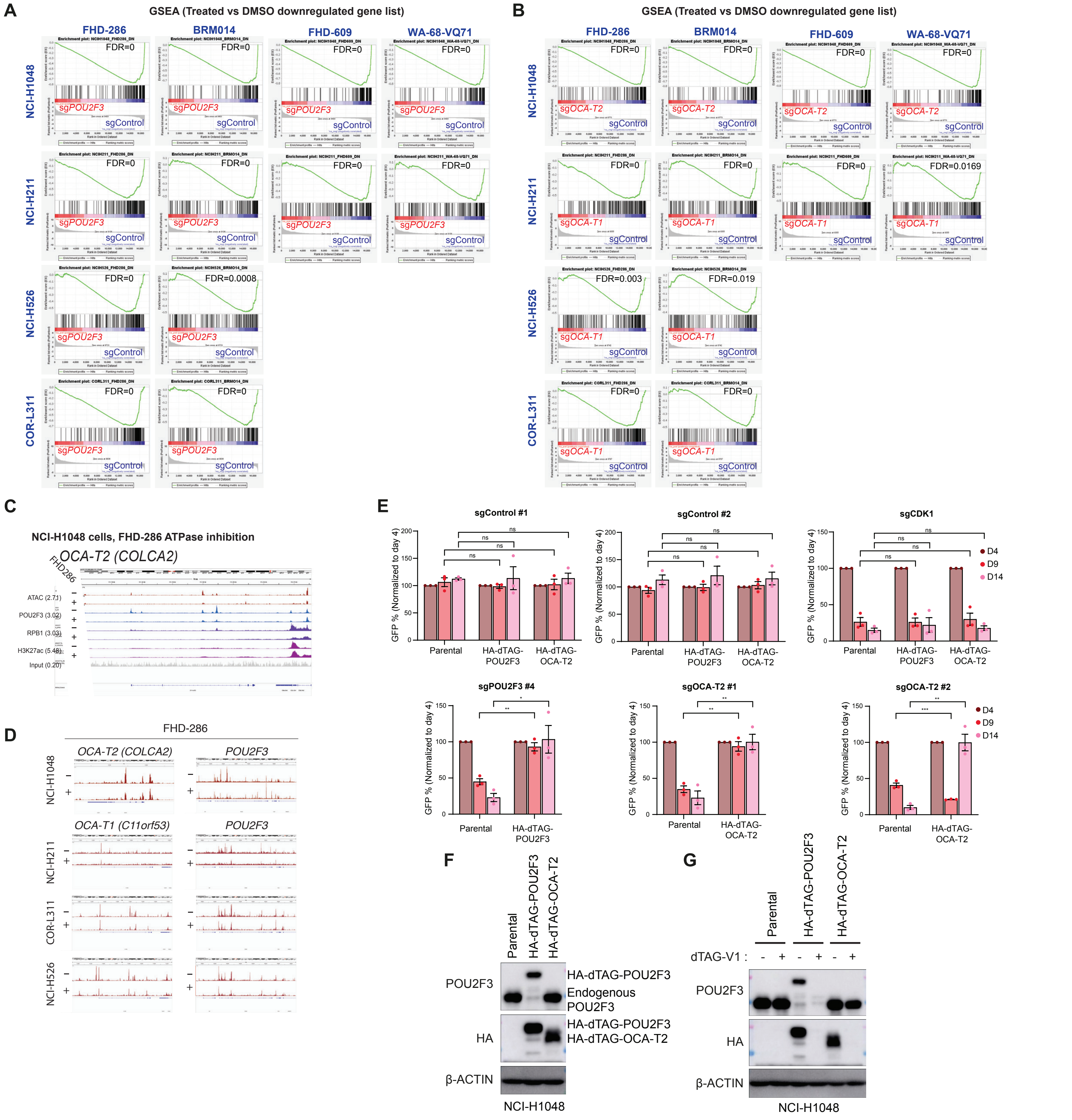


**Figure S6, Related to Figure 6. mSWI/SNF disruption-mediated chromatin and gene regulation at the POU2F3 TF and COLCA1/2 co-activator loci.** (**A-B**) GSEA enrichment plots (from Fig. 6A,C) of genes downregulated after treatment with the SMARCA4/2 inhibitors (FHD-286 or BRM014) or BRD9 degraders (FHD-609 or WA-68-VQ71) at 100 nM for 72 hours comparing sgPOU2F3 vs. sgControl (A) or sgOCA-T1 (for NCI-H211, NCI-H526, COR-L311) or sgOCA-T2 (for NCI-H1048) vs. sgControl (B) from Wu et al., 2022. n=3 biological replicates for FHD-286 and FHD-609. n=2 biological replicates for BRM014 and WA-68-VQ71. FDR q-values are indicated. (**C**) Chromatin targeting (ChIP-seq for POU2F3, RBP1, H3K27Ac) and accessibility regulation (ATAC-seq) at the OCA-T1(COLCA2) locus in NCI-H1048 cells treated with FHD-286. (**D**) ATAC-seq peaks over the POU2F3, OCA-T2 (COLCA2) and OCA-T1 (c11orf53) loci in DMSO and FHD-286 treatment conditions in the cell lines indicated. (**E**) Competition-based proliferation assays in NCI-H1048 Cas9 cell line co-transduced with the indicated cDNAs and sgRNAs (expressing GFP) to assess the functionality of the constructs. cDNAs were engineered to be resistant to Cas9/sgRNA-mediated cutting. Data are presented as mean +/- SEM normalized to percent GFP-positive at day 4 after infection. n=3 biological replicates. (**F**) Immunoblot analysis of NCI-H1048 Cas9 cells expressing exogenous sgRNA resistant HA-dTAG-POU2F3 with the knockout of endogenous POU2F3, or expressing sgRNA resistant exogenous HA-dTAG-OCA-T2 with the knockout of endogenous OCA-T2 via CRISPR, or parental NCI-H1048 cells. (**G**) Immunoblot analysis of cells shown in F after overnight treatment with dTAG-V1 at 100 nM.

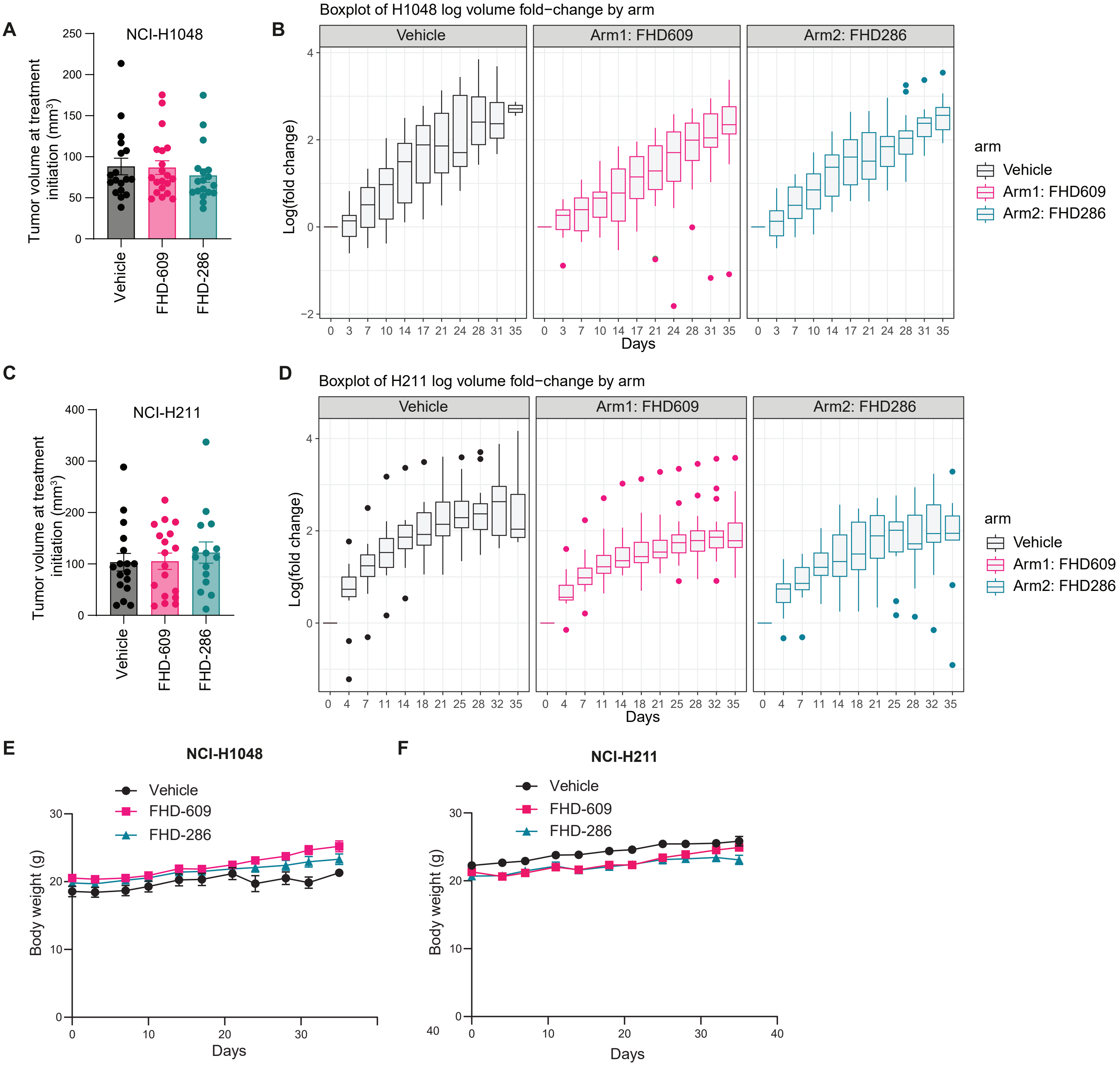


**Supplemental Figure 7. Related to Figure 7.** (**A,C**) Tumor volume in mm3 at treatment initiation for each treatment groups in NCI-H1048 (A) and NCI-H211 (C) xenografts. (**B,D**) Box plots of log fold-change in NCI-H1048 (B) or NCI-H211 (D) xenograft tumors treated with vehicle (Black), FHD-609 (0.5mg/kg IP QD) (pink) or FHD-286 (1.5mg/kg PO QD) (blue) for 35 days corresponding to the end of treatment. Data show log fold-change in tumor volume relative to tumor volume at treatment start (D0). For A,B; n=18 tumors from 9 independent mice (Vehicle), n=20 tumors from 10 independent mice (FHD-609), n=18 tumors from 9 independent mice (FHD-286). For C,D; n=17 tumors from 10 independent mice (Vehicle), n=18 tumors from 10 independent mice (FHD-609), n=15 tumors from 9 independent mice (FHD-286). (**E-F**) Body weights of mice enrolled in the NCI-H1048 (E) and NCI-H211 (F) xenograft efficacy treatment studies treated continuously daily for 35 days with vehicle (HP-β-CD), FHD-609 (0.5 mg/kg IP), or FHD-286 (1.5 mg/kg PO).
